# Cross Tissue DNAm Biomarker Prediction using Transfer Learning

**DOI:** 10.1101/2024.06.01.596949

**Authors:** Kristen M Mcgreevy, Brian H Chen, Steve Horvath, Donatello Telesca

**Affiliations:** Department of Biostatistics, University of California, Los Angeles, USA; San Francisco Coordinating Center, California Pacific Medical Center Research Institute, San Francisco, California, USA; Department of Epidemiology and Biostatistics, University of California San Francisco, San Francisco, California, USA; The Herbert Wertheim School of Public Health Human Longevity Science, USA; Altos Labs, UK

**Keywords:** DNA methylation, DNAm Biomarkers, cross tissue prediction, Transfer Learning

## Abstract

DNA methylation (DNAm) is an epigenetic mechanism vital for regulating gene expression and influencing disease states. Developing accurate DNAm biomarkers often requires data from specific tissues, which are sometimes difficult to access. This study explores the use of Transfer Learning (TL) to predict blood DNAm biomarkers using saliva DNAm data, aiming to overcome limitations posed by sample size and tissue accessibility. We developed TL-based algorithms that integrate DNAm data from multiple tissues. These algorithms were evaluated against traditional Lasso regression and direct saliva DNAm estimates. Our results show that TL significantly improves the prediction accuracy of DNAm biomarkers, outperforming traditional methods in 20 out of 26 biomarkers. We further validated our models using independent datasets, demonstrating that TL-derived predictions reflect known biological relationships, such as sex differences in telomere length and the impact of smoking on DNAm biomarkers. Our findings highlight the potential of TL in enhancing DNAm biomarker prediction across tissues, providing a valuable tool for epigenetic research. The developed algorithms and methodologies are accessible to researchers, fostering advancements in personalized medicine and aging research. This study establishes a framework for utilizing TL to bridge the gap between accessible and pertinent tissue data, paving the way for more accurate and versatile DNAm biomarker applications.

**ACM Reference Format:** Kristen M McGreevy, Brian H Chen, Steve Horvath, and Donatello Telesca. 2024. Cross Tissue DNAm Biomarker Prediction using Transfer Learning. 1, 1 (June 2024), 43 pages. https://doi.org/10.1145/nnnnnnn.nnnnnnn

## 1 INTRODUCTION

DNA methylation (DNAm) is an epigenetic mechanism that varies across tissues and contributes to cell type, regulates gene expression, and influences disease states [1]. Beyond this, DNAm has been leveraged for developing surrogate estimates, known as DNAm biomarkers, for a variety of phenotypes including age [2, 3], mortality risk [4], fitness [5], blood cell counts [6], and others [7–9]. DNAm continues to be studied offering a window into the biological processes and aging within cells [10]. Traditionally, blood has been the tissue of choice for DNAm biomarker development, serving as a versatile medium that interacts with and carries information from an array of organs. However, this choice is not without its limitations. The performance of DNAm based biomarkers is inherently tied to the relevance of the tissue that DNAm is measured in [11, 12], where biomarkers built with tissues more related to the condition or trait of interest are likely to be more accurate and informative.

Yet, herein lies a significant challenge—the tissues most pertinent to certain diseases or conditions, such as the brain, adipose, and bone, are often the hardest to access. Their collection is typically invasive, painful, and expensive, leading to small sample sizes. This scarcity significantly limits the development of tissue-specific DNAm biomarkers. While some research has looked at estimating methylation across tissues [13], algorithms that do this without having access to multiple tissue data are still unavailable. While some research has looked at estimating methylation across tissues [13] or predicting species’ average methylation across tissues [14], algorithms are not available for individual level prediction yet or without needing access to multiple tissue data. Moreover, simply measuring these tissues and applying current DNAm biomarkers may not yield meaningful insights, a point underscored by researchers who advise caution when using methylation markers from surrogate tissues [15].

This landscape delineates two explicit needs in the field of DNAm research. First, there is an urgent requirement for using more accessible tissues to accurately measure biomarkers of interest [11, 16]. Saliva emerges as a promising candidate, offering a non-invasive and easily accessible alternative to blood [17]. Its collection is patient-friendly and uncomplicated, allowing for larger sample sizes and broader applications. Second, there is a pressing need for novel methodologies that can develop accurate biomarkers in tissues traditionally challenged by inadequate sample sizes [16, 18].

Moreover, this context highlights an area for innovation: the use and understanding of Transfer Learning (TL) in the realm of DNAm research and biomarker development. TL, a powerful technique in machine learning, is adept at leveraging information from related data sources and applying knowledge from one context to enhance precision in another [19]. TL has been used in bioinformatics for tasks like imputing the methylome in low-coverage cases [20] and augmenting gene expression data [21], but its application in DNAm biomarkers, particularly for cross-tissue prediction, has yet to be explored.

Our study aims to address these needs from three angles. We demonstrate how transfer learning can improve DNAm biomarker accuracy, especially with limited data sources. We introduce cross tissue prediction algorithms for estimating common blood DNAm biomarkers using saliva DNAm. Additionally, our study offers practical tools and guidelines for researchers, empowering them to implement TL methods and develop algorithms tailored to their specific biomarker interests. Through these contributions, we aspire to provide easily usable methods for researchers to better estimate their favorite biomarkers across different tissues by leveraging shared information from other tissues, as well as methods to develop more accurate biomarkers in typically inaccessible tissues by combining data from similar sources.

Here, we adapt TL methods and develop cross tissue prediction algorithms. We focus on validating these algorithms with three key benchmarks deemed critical for any robust cross-tissue prediction algorithm. First, we assess whether TL offers an advantage in predicting blood DNAm biomarkers from saliva DNAm by comparing accuracy to direct saliva DNAm estimates. Second, we compare the TL algorithms against conventional Lasso regression to evaluate the benefits of adopting advanced computational techniques for developing cross tissue prediction algorithms. Third, we evaluate the ability of our algorithms to extend beyond mere accuracy and reflect biologically meaningful and established relationships in a variety of validation datasets.

## 2 METHODS

The core aim of our study was to assess the utility of Transfer Learning (TL) in the context of DNA methylation (DNAm) and DNAm-based biomarkers. Our analytical strategy is predominantly built upon the translasso framework [21]. In our research, we prioritized this class of algorithms due to their compatibility with the widely utilized Lasso regression techniques [22]. This choice ensures that our methods can be seamlessly integrated by researchers accustomed to Lasso regression, thereby promoting wider adoption within the field. Lasso, as opposed to elastic net regression, was chosen for simplicity and future methods can extend to other penalized regression frameworks. We employ a transfer learning paradigm for high-dimensional linear regression, utilizing prediction and estimation procedures outlined in the translasso study [21] to investigate whether transfer learning can be utilized in high dimensional epigenetics to improve DNAm biomarker prediction across tissues.

Our methods section has the following format. First, we describe the basic model set-up and datasets used. Then, we describe the transfer learning methodology that we employ. After that, we describe the variations to this methodology that we explore in order to optimize parameterization and TL application to our cross tissue DNAm setting. We then describe how we classify optimizations and develop the final cross tissue prediction algorithms. Finally, we describe the validation process for testing our algorithms in test datasets.

### 2.1 Subsetting Potential Covariates: C+S and C Method

We developed two distinct algorithms for predicting DNA methylation (DNAm) biomarkers across different tissues. The first algorithm, referred to as the C+S method, combines both the DNAm biomarkers derived from saliva and the raw methylation values from saliva samples. This approach allows for the saliva DNAm biomarkers to be ‘updated’ via methylation loci, enhancing their predictive accuracy. The terminology “C+S” references **C**pGs and **S**aliva DNAm Biomarkers being included as covariates. The second algorithm, known as the “C” method, exclusively uses saliva methylation values, ie CpGs, to predict blood DNAm biomarkers. This CpGs-only strategy is especially relevant when creating new biomarkers or when faced with datasets that lack a significant number of CpGs necessary for computing DNAm biomarkers, often due to variations in array platforms. By focusing solely on saliva methylation values, the C method provides a valuable tool in situations where comprehensive CpG data is unavailable or when developing novel biomarkers.

We restrict the potential covariates to predict each blood DNAm biomarker to CpG loci known to be informative to epigenetic clocks and conserved across commonly used methylation arrays. These loci are those published in public DNAm epigenetic clocks, specifically DNAmAge [2], DNAmAgeSkinBloodClock [23], DNAmHannumAge [3], DNAmPhenoAge [24], DNAmFitAge [5], Zhang [18], MethylDetectR [25], EpiTOC [26], and the PanMammalianClock [27]. Because some loci are not available across different array platforms, we included only CpG loci conserved across the 450K and EPIC array. This resulted in 6662 unique CpG loci to be included. For our C+S method, our potential covariates included 6663 variables: 6662 methylation sites and the saliva DNAm biomarker. In our paper, we refer to the saliva DNAm biomarker as the estimated DNAm biomarker if the current algorithms were applied directly to saliva methylation values, regardless if the algorithms were built for saliva or other tissues. For each cross tissue prediction algorithm, only 1 saliva DNAm biomarker will be included as a covariate. For the C method, we further reduced the methylation sites included to only those conserved across the 450K, EPIC, and Mammalian40K array. This resulted in 1307 unique CpG loci to be potential covariates in our C method. This is not only conducive to the development of human biomarkers but also holds potential for direct application to animal studies because the loci are conserved across mammalian species. For example, animal models measuring DNAm under calorie restriction (CR) could be integrated into our research to better inform the conserved epigenetic changes in humans from CR. Overall, developing and comparing algorithms in these two cases informs us of information loss or gain when implementing TL in DNAm contexts.

### 2.2 Datasets

Our “target” data includes datasets that have DNAm values in both saliva and blood. These datasets include the tissues of the targeted research objective: predicting blood DNAm biomarkers from saliva DNAm. We refer to “auxiliary” data as datasets that include DNAm in two tissues that are not saliva and blood. Finally, we include validation datasets, which are datasets not used in the TL algorithm development stage, but instead are for testing our developed algorithms in. All the datasets used have been previously described elsewhere; we provide brief summaries here.

#### 2.2.1 Target Data

Our target data consisted of 6 independent datasets that included DNAm in saliva and blood tissues for the same individuals. GSE111165 (n=33), GSE214901 (n=19), GSE159899 (n=19), GSE130153 (n=22), GSE59507 (n=4), and GSE73745 (n=12) for a total of 109 samples in the target datasets. In developing our methods, we employ a Leave-One-Data-Out (LODO) methodology that builds the TL model in 5 of the target datasets and tests in the 1 held out target dataset. This process is repeated for each target dataset, resulting in 6 LODO iterations per method, and the weighted average by target dataset size is used to calculate the final LODO estimate.

GSE111165 collected samples from blood, saliva, buccal, and brain tissue in epilepsy patients undergoing brain resection. DNAm was measured with both the 450K and EPIC arrays. GSE214901 measured brain, blood, saliva, and buccal in Japanese individuals undergoing neurosurgery, aged 13-73. DNA methylation was measured using the EPIC array. GSE159899 collected and measured methylation in whole blood, saliva, and T-cells using the EPIC array. GSE130153 measured saliva and blood sample methylation with the 450K array. GSE59507 measured multiple tissues from male crime scene samples aged 20-59 including blood, saliva, and semen. Methylation was measured using the 450K array. GSE73745 measured methylation in people with respiratory allergies and healthy controls in saliva and mononuclear blood cells. Methylation was measured using the 450K array.

(DT: Perhaps add some references to where these data sources have been published?)

#### 2.2.2 Auxiliary Data

Our auxiliary datasets consisted of 5 independent datasets that included DNAm measured in two different tissues for the same individuals. Comprehensive Assessment of Long-term Effects of Reducing Intake of Energy (CALERIE) included muscle and adipose (n=130), GSE111165 included buccal and blood (n=27), GSE214901 included buccal and brain (n=19), TwinsUK included adipose and skin (n=136), and GSE48472 included fat and blood (n=6) for a total of 318 samples in the auxiliary datasets. For each of these datasets, they are presented where the first tissue listed is the tissue used as the potential covariates, and the second tissue is the tissue used for the outcome. For example, in CALERIE, muscle DNAm is used in the *X* and adipose DNAm biomarkers are used in the *Y*. In the TwinsUK data, original person ID’s were not available for each tissue dataset, so individuals were matched across tissue based on SNPs, age, BMI, twin zygosity, and matching family IDs.

#### 2.2.3 Validation Datasets

We use four datasets that measure phenotypic variables known to relate to DNAm biomarkers and DNA methylation. This included GSE119078, GSE148000, GSE149747, and HorvathHIV. The first has 59 saliva samples measured in males (n=25) and females (n=34) with and without Celiac’s Disease on the 450K array. No differences were observed by disease group to the saliva DNAm biomarkers, and to our best knowledge, differential methylation from saliva DNAm for Celiac’s has not been observed elsewhere. This dataset is used to validate the relationship between sex and telomere length. The second dataset includes 26 asthma, COPD, and healthy patients where DNAm was measured in sputum using the 450K array, which is similar to, but not identically saliva. Age was observed to be different between the three disease classifications, so all models include age as a covariate. This dataset was used to look at differences in predicted DNAm biomarkers to disease status, relate predicted blood cell count models to measured lymphocyte percentage, and lung health DNAm biomarkers to reported cumulative smoking pack years. The third dataset is a exercise, diet, and sleep intervention study with saliva DNAm measured at baseline, 4 weeks, and 8 weeks after intervention on the EPIC v1 array. The HorvathHIV data measured methylation with a custom array (HorvathMammalMethylChip) from 661 samples across 11 human tissues (adipose, blood, bone marrow, heart, kidney, liver, lung, lymph node, muscle, spleen and pituitary gland) [12]. This sample included 133 clinically characterized, deceased individuals, including 75 infected with HIV. Based on the results of the initial study, we restricted our analysis to tissues with substantial sample sizes, relevant to saliva, and accessible- being lymph, muscle, and adipose.

When available, DNAm was processed using bioconductor minfi and SeSAMe packages in R with Noob normalization. This was not possible for GSE130153 which provides only processed DNAm data and not the original idat files. In addition, 3 datasets (GSE73745, 159899, and 130153) did not provide chronological age, which is required for some DNAm clocks. In these cases, their predicted age from DNAm was used in place of their chronological age. After pre-processing, DNAm data was uploaded to the online DNAm clock calculator to calculate DNAm biomarker values (https://dnamage.clockfoundation.org). The resulting epigenetic clocks were used as end points in our prediction models.

### 2.3 Transfer Learning Methodology

We are interested in estimating a set of regression coefficients *β*, mapping DNAm from saliva to DNAm biomarkers in blood. The model of interest is

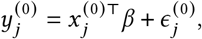

where *y*_*j*_ is the *j* th blood DNAm Biomarker, *x* is the vector of saliva DNAm inputs, and (0) refers to the target data. Because we develop two different algorithms for each DNAm biomarker, for the C+S method, *x* includes the *j* th saliva DNAm biomarker and saliva methylation values (1 × 6663 size vector), whereas for the C only method, *x* includes only saliva methylation beta values (1 × 1307 size vector).

Together with the target data, we observe additional data from a collection of auxiliary studies, denoted as *K*, to enhance our estimation of the vector *β*. Each auxiliary study provides samples that may vary in their relevance to the target data. The incorporation of these auxiliary samples into our analysis can take various forms—they can be merged into a single dataset, kept separate, or grouped into distinct combined datasets. To accommodate these different integration strategies, we define *L* as the ensemble of auxiliary data configurations utilized. Specifically, *L* represents any subset or combination of the *K* studies that we use for informing the estimation of *β*.

Mathematically, let us consider *K* auxiliary samples indexed by *k* = 1, … , *K* . The set *L* then represents a collection of these samples in various configurations, such as *L* = {(4) , (4, 1) , (4, 1, 3), (4, 1, 3, 2)}, or as a single combined set *L* = {(1, 2, 3, 4)}, or as individual datasets *L* = {(1), (2), (3), (4)} when *K* = 4. Here, the first instance illustrates *L* containing three groupings of *l* combining multiple auxiliary datasets (as is performed in the default settings of TransLasso), the second instance shows *L* as one grouping that combines all auxiliary datasets, and the third instance depicts *L* with each *l* representing an individual auxiliary dataset. For scenarios where auxiliary samples are considered individually, such as in the Oracle 1df method, we can substitute *l* with *k* in our notation. This substitution reflects the scenario where each auxiliary sample is assessed separately to inform the model.

The general methodology can be summarized into five primary steps below.

**Step 1:** Perform Lasso on the Target Data (saliva DNAm →blood DNAm biomarker):

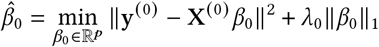

**Step 2A*:** Determine the informative set, *L*, from auxiliary data, *K* , as detailed in Section 2.3.1

**Step 2B:** Perform Lasso on Informative Auxiliary Data for each *l* ∈ *L* (any tissue DNAm → any other tissue DNAm biomarker):

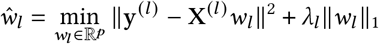

**Step 3:** Perform Lasso on Target Data Residuals (saliva DNAm→ Residual blood DNAm biomarker):

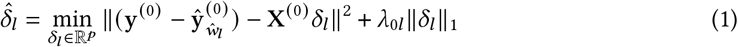

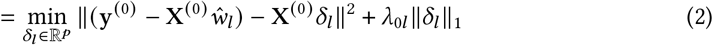

**Step 4:** Calculate coefficients estimated from the auxiliary samples with or without a threshold for combining:

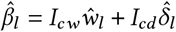

**Step 5:** Combine 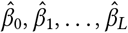 for ∀*m* ∈ 0 : *L* using the squared prediction error on the target data.

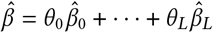

where

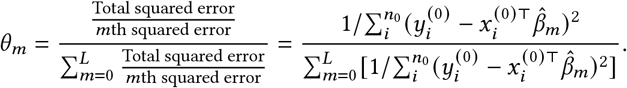

In Steps 1-3, the tuning parameters *λ*_0_, *λ*_*l*_, *λ*_0*l*_ are selected through cross validation and are described in greater detail in the next Methods section. In Step 4, *I*_*cw*_ and *I*_*cd*_ are indicator vectors that indicate which coefficients of *ŵ*_*l*_ and 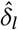 are of magnitude greater than or equal to the threshold value of *c*. This is also in the next Methods section, where we detail the different constants we use, including allowing any coefficient magnitudes to be added together. In the weighting scheme *θ*_*l*_ of step 5, the error is calculated in the training target data. The term, 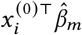 is the the prediction for individual *i* in the target data using model *m*, and the summation over *i* aggregates squared prediction errors over all individuals. The weights are based on the inverse of the squared error for each model’s predictions on the target data, which gives more importance to models with lower errors. We note that this is *not* the error from the left out target data, because then this algorithm would not be applicable to researchers developing new algorithms without validation data. Error from LODO is used at a later step, however, to combine the coefficients from our various TL models in a Super Learner process to obtain the final coefficients.

This methodology is described with all 6 target datasets together in y^(0)^ and X^(0)^ , which is the case for our final algorithms. However, in the process of testing parameterization and TL method specification, we employ a Leave-One-Data-Out (LODO) process, meaning 1 target dataset is held out and the remaining 5 are used to develop the TL coefficients in the steps described above. As such, for developing the TL algorithm for each biomarker and each tested TL method, the 5 steps were performed 6 times to capture *j* = 6 LODO folds. This results in 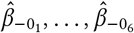 for each *j* th fold (total of 6 target datasets), which are the target data coefficients when the *j* th target data is left out 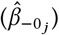 . After each fold is run, the 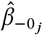 coefficients are applied in the *j*th dataset to calculate the LODO error and correlation for comparing the TL methods. The LODO error and correlation are a sum of the metrics weighted by the left out dataset size, *n*_*j*_. For example, the LODO total squared error is calculated as:

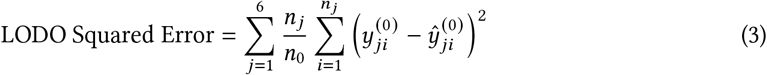

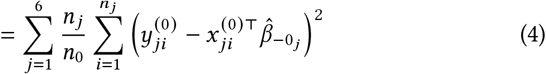

where 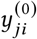 is the blood DNAm biomarker of interest for the *i*th person in the *j*th left out target dataset and 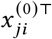 is vector of saliva DNAm for the *i*th person in the *j*th left out target dataset.

#### 2.3.1 Calculating Informative Auxiliary Sets (Step 2A)

In Step 2A, we use * to denote the possibility to skip this step, as is the case for the Oracle methods, where informativeness is a-priori known and therefore does not need calculated. We employ two oracle methods. One, which we refer to as Oracle A0, treats all auxiliary datasets as equally informative, and *L* is the combined set of all *K* auxiliary data. In the second oracle method, deemed Oracle 1df, each auxiliary dataset is considered informative, but not necessarily equal. In this case, *L* is exactly *K* , with each auxiliary dataset being treated individually.

When we are determining the informativeness of auxiliary datasets, we construct *L*, the set of informative auxiliary datasets. In summary, this algorithm computes differences in marginal correlations between auxiliary datasets and the target datasets. The most significant of these differences calculates an index 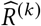 that gives an indication of the informativeness of the *k*th auxiliary dataset. These are ranked and auxiliary datasets are taken in sequence to produce the informative set *L*.

**Step 1:** Calculate 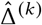:

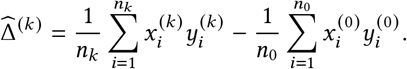

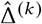 holds the marginal statistics for the *k*^*th*^ auxiliary dataset, which is the average difference between the *k*^*th*^ auxiliary dataset CpG’s and the target dataset CpG’s correlation to the DNAm biomarker outcome, *y*. The term *X*^′^*y* captures how each predictor variable individually correlates with the DNAm biomarker variable. Thinking of *y* as a vector in a space, and each column of *X* as another vector in that space, then each component of *X*^′^*y* is essentially the projection of *y* onto each of those predictor vectors. In this framework, we are taking the average difference of the projection between the auxiliary data and target data for each . (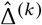 is a p × 1 vector)

**Step 2:** Pick out the most significant components of 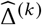 by obtaining the top 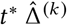 (largest differences) between the auxiliary and target data, called 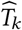. As such, 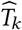 are indices of the largest marginal statistics for each auxiliary dataset. The default value for *t*\* is one third the target dataset size (*n*_0_ /3), and we also explore variations to include more 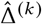. This parameter changes how many of the largest differences are considered. Call this subset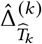.

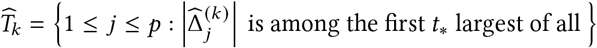

By taking the largest values, we are capturing the most unique differences between the auxiliary and target data. Capturing the largest differences allows us to understand how different the auxiliary data are from the target data.

**Step 3:** Calculate Sparse Index 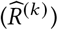 for the *k*^*th*^ auxiliary dataset.

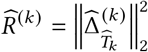

This estimated sparse index is the squared Euclidean norm of the vector 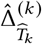. Essentially, this is summing the total squared differences of the top *t*^*^ correlation differences. Auxiliary datasets with smaller 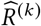 are more informative because the total deviation between the auxiliary and target data is small. This metric will be larger for auxiliary datasets that have more significant differences from the target datasets. After this, we use these sparsity indices to make candidate sets / subsets of the auxiliary data. We rank each 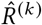 and take the smallest values in sequence to make *L* subsets of size 1, … , *K* . Specifically, the *l*^*th*^ candidate set is:

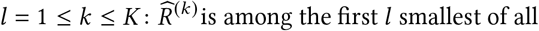

For illustration, we have 5 auxiliary datasets, and suppose the datasets’ rank of 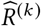’s is 4,1,2,3,5. Then *L* would be the 5 element set {(4) , (4, 1) , (4, 1, 2) , (4, 1, 2, 3) , (4, 1, 2, 3, 5)} . Computation proceeds as described above with each subset of auxiliary datasets being run together in the Lasso algorithm. In total, there will be *K* subsets each with 1, … , *K* auxiliary datasets included. Therefore, the most informative auxiliary dataset will be present in all *L* subsets.

### 2.4 Data Distinctions to Classical Transfer Learning

Our study falls within a unique category of ‘multi-source, multi-target, high-dimensional, homogeneous transfer learning’, however our data has novel characteristics not typically observed or studied in TL.

While conventional transfer learning might focus on domains with significant overlap or similarity, our approach blends data across various tissue types, some of which might share more characteristics than others. This heterogeneity is distinct in that it involves not only different tissues (with tissue specific methylation patterns) but also different relationships across two different, distal tissues. By predicting across tissues and incorporating data from different cross tissue predictions, we are seeking to capture universal within-person shared tissue signals. Furthermore, our target datasets, focusing on saliva DNAm predicting blood DNAm biomarkers, are aggregated from six different studies. This multi-study integration is uncommon in transfer learning, where target data is usually drawn from a single or more uniform source [**? ?** ]. While Horvath’s DNAmAge clock was built with multiple tissues, most other DNAm biomarkers are not [2]. The diversity in our target datasets adds complexity due to variations in study design, data collection methods, and participant characteristics. This methodology requires sophisticated data handling and model optimization to manage the heterogeneity across sources and domains. We explore the optimization of TL techniques for complex epigenetic problems with methods simple enough that other researchers can readily adopt.

### 2.5 Optimizing Transfer Learning Methodology for Cross-Tissue DNAm Prediction

The core aim of our study was to assess the utility of Transfer Learning (TL) in the context of DNA methylation (DNAm) and DNAm-based biomarkers. We made methodological modifications within the TransLasso framework to tailor the TL process for DNAm based biomarkers. The variations explored include lambda penalty optimization, coefficient thresholding, and the integration of auxiliary dataset information, which are delineated below.

#### 2.5.1 Lambda Penalty

The *λ* penalty in penalized regression models is typically determined through cross-validation (CV), either by identifying the *λ* that minimizes CV error or by selecting a more conservative *λ* within one standard error above the minimum. In transfer learning contexts involving multiple auxiliary datasets, optimizing *λ* for each dataset can be computationally intensive. We expand upon the TransLasso algorithm, which uses a constant calculated from the target dataset, *c*_0_, to define *λ*_*k*_ for each auxiliary set: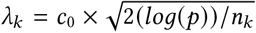. Our investigation included this approach and allowed for *λ* selection through CV in each auxiliary set, aspiring to harness dataset-specific nuances to inform *λ* choice.

In our CV *λ* selection process, *λ*_0_ is the optimal *λ* for the target dataset from CV on y^(0)^ and X^(0)^ , *λ*_*k*_ is the optimal tuning parameter for *ŵ*_*k*_ from y^(*k*)^ and X^(*k*)^ , and *λ*_0*k*_ is the optimal tuning parameter *λ*_*k*_ scaled by the target dataset size, *n*_0_. We scrutinized the constant *λ* and individualized *λ* approaches under the min and 1se parameterizations to determine the most suitable for our DNAm biomarker prediction. This approach was geared towards determining the optimal *λ* for our prediction problem, with the intent to refine the model’s predictive accuracy and reliability.

#### 2.5.2 Coefficient Thresholding

In the transfer learning process, a two-step coefficient computation is used for estimating coefficients from auxiliary data: initially within the auxiliary dataset to predict the outcome of interest (weights, *w*_*k*_), followed by an adjustment for biases between the target and auxiliary datasets (*δ*_*k*_). Lasso regression shrinks coefficients toward zero, which can result in some coefficients being small and near zero. To mitigate the incorporation of negligible coefficients from auxiliary sources, thresholding can be applied prior to combining the auxiliary weights and bias coefficients. While the conventional TransLasso method maintains coefficients exceeding a threshold of *λ* (described above), our exploration incorporated more lenient thresholds, including halving the threshold (0.5 × *λ*) and considering all coefficients, regardless of coefficient magnitude. We compare these three thresholding approaches and refer to them as ‘lambda’, ‘half lambda’, and ‘all’. This approach acknowledges the inherent characteristics of DNA methylation (DNAm) where the effects can be minuscule. By lowering or removing the thresholding, we may account for the subtlety of DNAm effects and improve the application of TL to DNAm data.

#### 2.5.3 Auxiliary Dataset Information

The relevance of auxiliary datasets to your target data and objective can often be ambiguous. An ‘oracle’ scenario would entail using only known informative auxiliary datasets; however, this is not always practical. Consequently, we need a mechanism to gauge the informativeness of auxiliary datasets. We examined both the oracle and estimation approaches for auxiliary dataset incorporation. In the oracle scenario, we considered all auxiliary data as either a single collective sample or as separate individual datasets, referred to as ‘Oracle A0’ and ‘Oracle 1df’, respectively. Informativeness of auxiliary datasets were computed as described above using the differences in marginal correlations. We varied the number of considered correlations to calculate auxiliary data similarity starting from the default value of one-third of the target dataset size (*n*_0_/3). For the C+S method, we considered the top 100, 500, 2000, and 6663 (all) correlation differences. For the C method, limited to the 40K array-conserved sites, we considered the top 100, 500, and 1307 (all) correlation differences. These methods are referred to as Rhat and the corresponding number of correlations considered, like ‘Rhat100’.

### 2.6 Evaluating TL Methods and Developing Final Algorithms

#### 2.6.1 Evaluating Transfer Learning (TL) Methodologies

To assess the operating characteristics of various TL algorithms, we conducted a comprehensive evaluation of their performance using Leave-One-Dataset-Out (LODO) correlation, mean squared error (MSE), and mean absolute percent error (MAPE). The results of this analysis, delineated by algorithmic variation, are presented in Table 2. This approach provides a holistic view of how different parameters and methods of integrating auxiliary data affect the predictive accuracy for DNA methylation (DNAm) biomarkers. It should be noted, however, that while this evaluation offers a broad understanding of algorithmic performance, it does not specifically address variations across individual DNAm biomarkers.

**Table 1.** Dataset Summary.

| Purpose | Dataset | N | Tissues |
| --- | --- | --- | --- |
| Target | GSE111165 | 33 | saliva, blood |
|  | GSE214901 | 19 |  |
|  | GSE159899 | 19 |  |
|  | GSE130153 | 22 |  |
|  | GSE59507 | 4 |  |
|  | GSE73745 | 12 |  |
| Auxiliary | CALERIE | 130 | muscle, adipose |
|  | GSE111165 | 27 | buccal, blood |
|  | GSE214901 | 19 | buccal, brain |
|  | TwinsUK | 136 | adipose, skin |
|  | GSE48472 | 6 | adipose, blood |
| Validation | GSE119078 | 59 | saliva |
|  | GSE148000 | 26 | sputum |
|  | GSE149747 | 122 | saliva |
|  | HorvathHIV | 28-58 | lymph, adipose, muscle, blood |

**Table 2.**
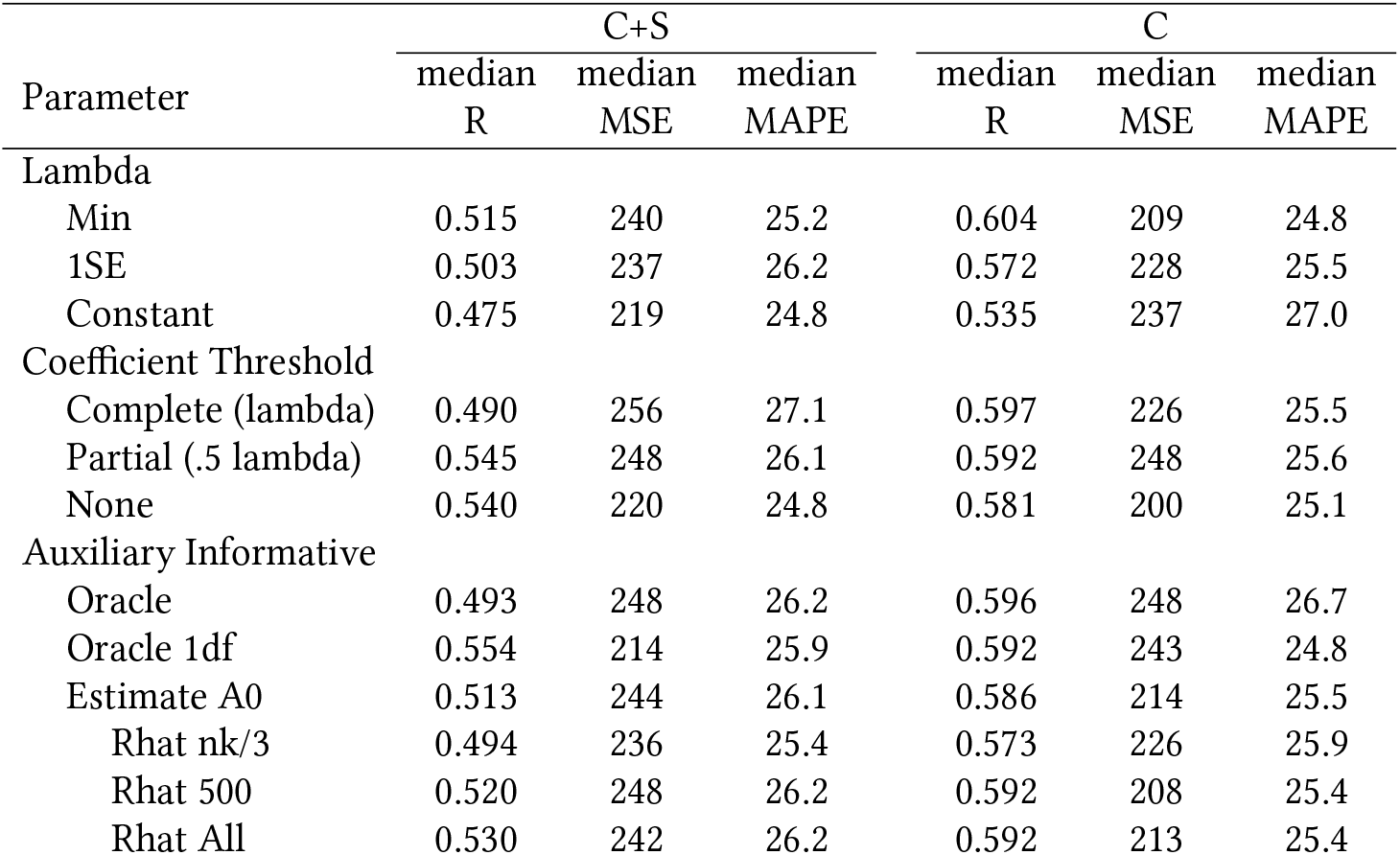
Optimal Parameters for Transfer Learning with DNAm.

| Parameter | C+S |  |  | C |  |  |
| --- | --- | --- | --- | --- | --- | --- |
|  | median<br>R | median<br>MSE | median<br>MAPE | median<br>R | median<br>MSE | median<br>MAPE |
| Lambda |  |  |  |  |  |  |
| Min | 0.515 | 240 | 25.2 | 0.604 | 209 | 24.8 |
| 1SE | 0.503 | 237 | 26.2 | 0.572 | 228 | 25.5 |
| Constant | 0.475 | 219 | 24.8 | 0.535 | 237 | 27.0 |
| Coefficient Threshold |  |  |  |  |  |  |
| Complete (lambda) | 0.490 | 256 | 27.1 | 0.597 | 226 | 25.5 |
| Partial (.5 lambda) | 0.545 | 248 | 26.1 | 0.592 | 248 | 25.6 |
| None | 0.540 | 220 | 24.8 | 0.581 | 200 | 25.1 |
| Auxiliary Informative |  |  |  |  |  |  |
| Oracle | 0.493 | 248 | 26.2 | 0.596 | 248 | 26.7 |
| Oracle 1df | 0.554 | 214 | 25.9 | 0.592 | 243 | 24.8 |
| Estimate A0 | 0.513 | 244 | 26.1 | 0.586 | 214 | 25.5 |
| Rhat nk/3 | 0.494 | 236 | 25.4 | 0.573 | 226 | 25.9 |
| Rhat 500 | 0.520 | 248 | 26.2 | 0.592 | 208 | 25.4 |
| Rhat All | 0.530 | 242 | 26.2 | 0.592 | 213 | 25.4 |

#### 2.6.2 Development of Optimized TL Algorithms

Our methodology for refining TL algorithms involved a detailed analysis of their performance across each DNAm biomarker. We calculated and then ranked both the correlation and prediction error for each TL method within individual DNAm biomarkers. This ranking was based on a weighted LODO MSE and the correlation between the predicted and actual blood DNAm biomarkers, with weights assigned in proportion to the size of the dataset left out. The most effective methods were identified based on their mean performance in terms of correlation and MSE across all DNAm biomarkers. The four top-ranked methods were subsequently applied to the entire target dataset, with the resulting coefficients being recorded.

In the final stage of our analysis, we employed a Super Learner approach to combine coefficients from these four top performing methods, thereby deriving the final algorithms’ coefficients for each DNAm biomarker. The weights assigned to each coefficient set were inversely proportional to the squared errors from the LODO analysis of the respective method. We opted for the Super Learner framework over the single best-performing TL method because of the former’s proven superiority in enhancing predictive accuracy and providing more stable estimates in regression models, particularly when multiple a priori techniques are utilized [28]. We developed the optimal TL algorithm separately for the two algorithms desired: one incorporating saliva DNAm biomarkers with 6662 CpGs as potential covariates and the other using only the 1307 CpGs conserved on the 40K array. Consequently, for each DNAm biomarker, we developed two cross-tissue prediction algorithms — one incorporating saliva DNAm biomarkers and the other based solely on saliva methylation beta values. However, as described in the section below, we do not provide all DNAm biomarker algorithms to researchers because the TL method does not always adequately predict DNAm biomarkers.

### 2.7 Validation of Algorithms

In our study, we embarked on a comprehensive evaluation of Transfer Learning (TL) to determine its potential contributions in the realm of cross-tissue prediction, particularly in the context of predicting blood DNA methylation (DNAm) biomarkers. Our evaluation framework was multi-faceted, focusing on three key benchmarks deemed critical for any robust cross-tissue prediction algorithm.

#### 2.7.1 Comparison of TL with Direct Saliva DNAm Estimates

Initially, we assessed whether TL offers an advantage in predicting blood DNAm biomarkers from saliva DNAm. This involved contrasting the efficacy of our TL algorithms with the baseline approach of directly computing biomarkers from saliva DNAm. The TL algorithms can be considered advantageous if they demonstrate superior performance in approximating blood DNAm biomarkers compared to using saliva DNAm as a direct surrogate.

#### 2.7.2 Benchmarking TL against Lasso Regression

We then compared the TL algorithms against conventional Lasso regression to evaluate the benefits of adopting advanced computational techniques. Lasso regression was applied solely to the target data, serving as a comparative baseline. In contrast, our TL methods leveraged both the target data and additional auxiliary cross-tissue DNAm data. The key distinction between the TL and Lasso methods lies in the incorporation of this auxiliary cross-tissue DNAm data in TL as both TL and Lasso used the same target and held out datasets for development and comparison. We analyzed and compared the Leave-One-Dataset-Out (LODO) correlations and errors generated by both the TL and Lasso algorithms. The TL algorithms can be considered advantageous for DNAm biomarkers if they demonstrate better prediction accuracy compared to Lasso techniques.

#### 2.7.3 Comparing TL Algorithms with Different Domains

Lastly, we scrutinized the differential predictive power and accuracy between our two distinct TL algorithms developed for each DNAm biomarker. One algorithm incorporated saliva DNAm biomarkers and beta values across 6662 CpG loci, while the other was confined to 1307 CpG sites alone. This comparative analysis aimed to determine whether the inclusion of saliva DNAm biomarkers as covariates significantly enhances predictive accuracy or if their inclusion is largely redundant in the context of these TL models.

### 2.8 Scope of Algorithms: Application to Validation Sets

The utility of these algorithms extends beyond mere accuracy; for them to resonate with and be adopted by the broader research community, they must demonstrate an ability to reflect biologically meaningful relationships. We explore this by examining if our predicted DNAm biomarkers have the same relationships that have been established between blood DNAm biomarkers and phenotypic variables across three distinct validation datasets. Then, we examine the robustness of input tissue type to our algorithms using a validation dataset with multiple tissues and HIV status.

We calculate the association between our predicted blood DNAmTL biomarker and sex to evaluate if our predictions align with known biological trends, where females tend to have longer telomere length compared to men of the same chronological age [29]. Next, we compare predicted blood DNAm biomarkers among people with COPD, asthma, and healthy controls, using sputum DNAm and adjusting for age. Research has demonstrated older DNAm age estimates in people with COPD and asthma compared to controls [30]. We also investigate the congruence of predicted blood cell count DNAm biomarkers with lymphocyte percentages. The study also reports person-reported cumulative (or lifetime) smoking pack years, and we evaluate the association to predicted DNAm biomarkers surrounding lung health and fitness: DNAmPackYears, DNAmFEV1, and DNAmVO2max blood biomarkers. Finally, we explore if our predicted blood DNAm biomarkers change in the expected direction from a longitudinal exercise, diet, and sleep intervention study. Here, we employ a main effects mixed model, controlling for age and study duration, to determine if our fitness-related blood biomarkers show expected improvements in the intervention group. This analysis also offers the unique opportunity to assess whether the newly developed blood DNAm fitness biomarkers exhibit expected improvements from an exercise intervention, an evaluation that has not yet been undertaken. These comparisons validate the predictive potential of the TL algorithms and demonstrate their robustness to application in novel data sources.

#### 2.8.1 Assessing Algorithmic Flexibility to Tissue Type

Because our TL approach incorporates information from multiple human tissues, we are interested in understanding if the algorithm is robust to tissue type. For example, instead of saliva DNAm, can we provide lymph nodes or other tissue DNAm and still have accurate blood DNAm biomarker predictions? We applied the C algorithms to three accessible tissues (lymph nodes, adipose, and muscle DNAm) in HIV-negative and HIV-positive individuals and compared the predicted DNAm biomarkers. This helps us assess if the predictions accurately reflect the accelerated aging and immune dysregulation associated with HIV infection.

### 2.9 Summary of Terminology Used

- **LODO**: Leave-One-Data-Out, meaning leaving 1 target dataset out at a time.
- **TL**: Transfer Learning
- **C+S Method**: One algorithmic set up which includes 6662 saliva CpG loci and the saliva DNAm biomarker as covariates to predict the blood DNAm biomarker.
- **C Method**: The second algorithmic set up which includes 1307 CpG loci as covariates to predict the blood DNAm biomarker.
- **Target Data**: Datasets that include saliva and blood DNAm from the same individual. There are 6 of these and are used to develop the algorithms. Referred to with ^(0)^ .
- **Auxiliary Data**: Datasets that include two tissues that are not saliva and blood from the same individual. There are *K* = 5 of these and are used to develop the algorithms. Referred to with ^(*k*)^ .
- **Validation Data**: Four datasets that are not used to develop the algorithms, but are used to apply the algorithms to validate the signatures.
- **Oracle A0**: TL method where all auxiliary datasets are a-priori specified as equally informative and combined into 1 auxiliary dataset
- **Oracle 1df**: TL method where all auxiliary datasets are specified as informative and used as individual auxiliary datasets
- **Informative Auxiliary Sets**: When not using the Oracle TL methods, we estimate which auxiliary datasets are informative for our target data. Referred to with *l* and *L*.
- **Min / 1SE**: The *λ* penalty term in Lasso that either minimizes the CV error or is the value that is at most 1 standard error above the minimum CV error.
- **lambda, half lambda, all**: The three different coefficient thresholds used for combining weights, *ŵ*_*k*_ , and bias coefficients, 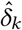, from the auxiliary data
- **Rhat**: The different number of correlation differences considered when determining auxiliary informative sets.

## 3 RESULTS

### 3.1 Optimal TL Algorithm

#### 3.1.1 Optimal Parameters for C+S Method

The best median LOOCV correlation was observed using the minimum lambda in the C+S methods (0.515) (Table 2). Interestingly, despite a relatively lower correlation of 0.475 under the constant lambda, determined based on CV in target data and auxiliary dataset size, the error metrics are slightly reduced. This suggests improved prediction accuracy despite a less strong linear relationship between the predicted and true blood DNAm biomarker. Settings without thresholding or partial thresholding results in comparable correlation with no thresholding having 1.3% lower error. Both of these methods outperform complete (lambda) thresholding with a difference of 0.05 correlation and 2.3% higher error, suggesting complete thresholding underfits the model by removing small, but informative coefficients.

The integration of auxiliary datasets through the Oracle 1df approach yielded the best correlation and the lowest Mean Squared Error (MSE) on average, with a median Mean Absolute Percentage Error (MAPE) similar to other methods incorporating auxiliary information. Contrary to expectations, the Oracle 1df method outperformed the standard Oracle approach, which treats all auxiliary datasets as a single combined source, improving correlation by 0.06 and maintaining comparable MAPE. Furthermore, the estimation of the informative set indicated a beneficial trend in incorporating more Rhats, as evidenced by a gradual improvement in correlation (R = 0.494, 0.520, 0.530) with negligible variation in error metrics. To understand variability in performance for the C+S method based on parameterization, we provide Supplemental Figure 2 for each biomarker. In summary, the C+S method’s optimal generalization employs the Oracle 1df approach, does not perform coefficient thresholding, and uses the minimal lambda derived from cross-validation in each dataset.

**Figure 1.**
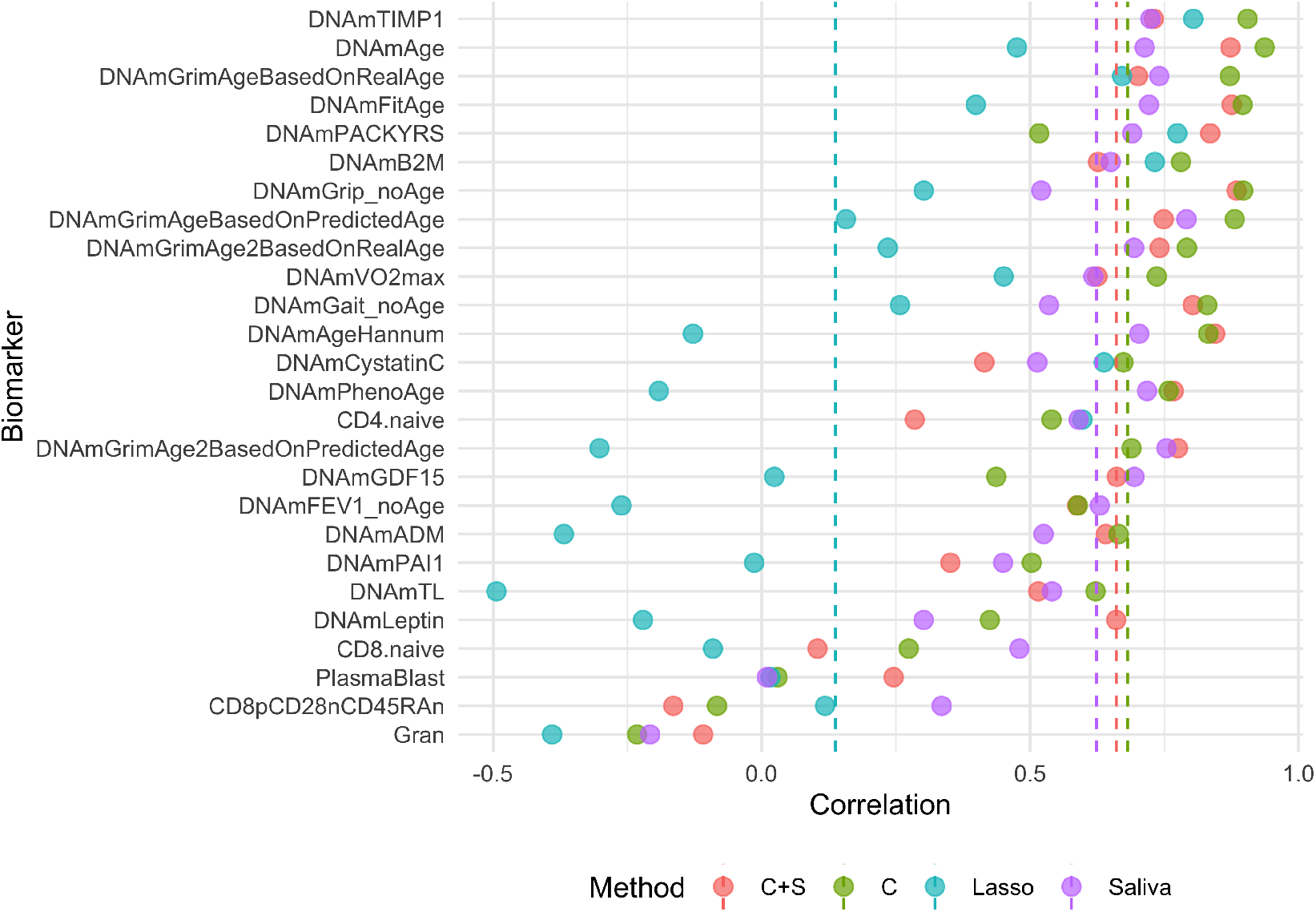
Correlation between True and Estimated Blood DNAm Biomarkers by top Performing TL Methods, Lasso, and Saliva Surrogates. Median correlation presented as dotted line. LODO Correlation presented for C+S, C, and Lasso methods.

**Figure 2.**
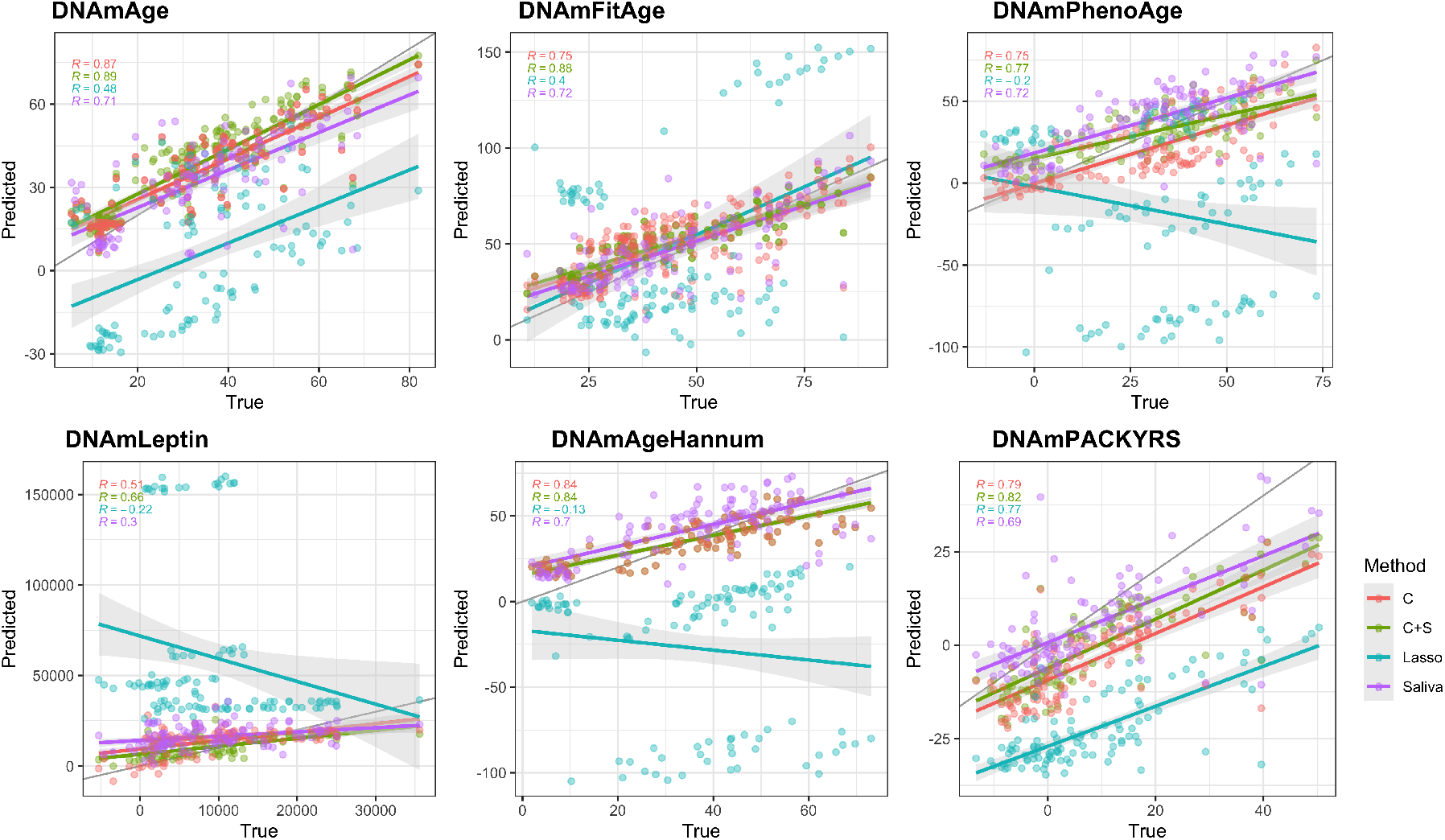
Scatterplots between True and Estimated Blood DNAm Biomarkers. LODO Correlation presented for C+S, C, and Lasso methods.

#### 3.1.2 Optimal Parameters for C Method

In contrast, the C method consistently demonstrates higher median R values across most parameters compared to C+S. This suggests a stronger correlation with the actual blood DNAm biomarker under the C setting. Echoing the C+S method’s findings, the minimum lambda from auxiliary data CV was preferred, achieving the highest average R of 0.604 and lowest MAPE of 24.8% (Table 2).

In a departure from C+S, the C method favored more rigorous coefficient thresholding. Specifically, the median R of 0.597 with complete thresholding was 0.016 higher than settings without thresholding. This divergence between methods suggests a greater number of CpGs are needed to estimate the blood biomarker in the absence of original tissue DNAm biomarkers (C method), however, this also propagates noise in the prediction, which can be corrected by removing smaller coefficients. Finally, auxiliary methods using either Oracle 1df or estimating the informative set with more Rhats perform comparably well based on R and MAPE. While Oracle and Oracle 1df have similar correlations, Oracle 1df decreases error by approximately 2%.

Thus, the C method’s best practice includes the minimal CV lambda from each auxiliary dataset combined with complete (lambda) coefficient thresholding. Using either Oracle 1df or estimating the informative sets with more Rhats are similarly efficacious.

#### 3.1.3 General Recommendations

Both C+S and C methods favor the Oracle 1df method and the minimal lambda. They diverge on coefficient thresholding — more thresholding benefits the C method, while less thresholding benefits the C+S method. These recommendations, however, do not account for individual biomarker variability, which may influence the optimal choice for a specific biomarker. Researchers with the resources to conduct an LOO procedure are advised to employ the TL tuning function to determine the best parameters for their specific scenario. In the absence of such a procedure, our broad recommendations provide a starting point.

### 3.2 Final Algorithms

Our previous section evaluated individual parameters, however, it’s crucial to recognize that the best individual parameterizations might not synergize when combined. Therefore, we ranked each TL method within each biomarker to determine the most efficacious models. These results generally aligned with our recommendations, particularly for the C+S method where the Oracle 1 df, min lambda, and all coefficient setup emerged as the top-ranked for lowest Mean Squared Error (MSE) and highest correlation among all C+S TL methods. Despite its superior performance, it didn’t achieve a universal top rank, highlighting the presence of multiple algorithms with nearly identical performance across various methods. The top 4 C+S methods were Oracle A0 1df, min lambda, all coef; Oracle A0, 1se lambda, half coef; Estimate A0 Rhat nk/3, 1se, half coef; and Estimate A0 Rhat All, 1se lambda, half coef. For the C method, the top 4 algorithms were OracleA0 1df, 1se lambda, all coef; Estimate A0 Rhat 500, min lambda, all coef; Estimate A0 Rhat nk/3, min lambda, lamb coef; and Oracle A0, min lambda, lambda coef. The variation in top methods reaffirms that no single TL method suits all scenarios. Given the absence of one-size-fits-all solution, we used a Super Learner approach to generate the final coefficients for each biomarker. This strategy capitalizes on the strengths of each top algorithm, mitigating their individual weaknesses and catering to the specificities of each biomarker. The final algorithms for each biomarker are readily accessible on our GitHub repository at https://github.com/kristenmcgreevy/EpigenTL, and are also provided in the Appendix section 7.

### 3.3 Algorithmic Comparison

#### 3.3.1 Comparison to Saliva Surrogates

In the comparative analysis of Transfer Learning (TL) algorithms for estimating blood DNA methylation (DNAm) biomarkers from saliva DNAm, we found that 20 out of 26 biomarkers were more accurately predicted by TL methods than by their saliva DNAm surrogates, based on Mean Squared Error (MSE) and correlation metrics. When separating the comparison between the C+S method and C method to saliva, 18 and 15 biomarkers were more accurately predicted with the TL methods, respectively. The amount of improvement using either of the Transfer Learning methods varied by biomarker. Some biomarkers saw drastic improvement, such as DNAmGrip_noAge and DNAmLeptin, which had 0.364 and 0.364 stronger correlation, respectively. DNAmLeptin also saw a 7 figure improvement in MSE when using our C+S algorithm. For our provided algorithms, the average LODO correlation improvement of our algorithms compared to the saliva surrogate itself is 0.120. This improvement is not universal across all DNAm biomarkers, however. One biomarker—Gran—showed notably poor correlations across TL, Lasso, and saliva surrogate methods, prompting its exclusion from any saliva to blood biomarker recommendations. In addition, CD8pCD28nCD45RAn did not have good performance with C+S method, C method, or Lasso, however the saliva surrogate had a decent correlation (0.33) (Supplemental Table 1).

#### 3.3.2 Comparison between TL and Lasso

When comparing TL algorithms directly against Lasso regression, TL algorithms outperformed Lasso for 23 out of 26 biomarkers. This underscores the significant potential of TL algorithms in the generation of new DNAm biomarkers. Additionally, the Lasso algorithm averaged the lowest/worst rank in MSE and Correlation across all TL and Saliva surrogate methods within each biomarker. Even when Lasso outperformed the TL methods, it did not outperform the saliva surrogate alone. With these results, we did not observe any instance where Lasso would be more beneficial than TL when saliva surrogates are available. As such, we strongly encourage researchers to adopt TL methods in place of Lasso when similar data are available, like when different tissue DNAm samples are available. Additional metrics evaluating median absolute percent error are presented in the Appendix Section 1.

#### 3.3.3 Comparison of C+S to C method

Unexpectedly, 9 biomarkers estimated with the C method surpassed the performance of the C+S method. This suggests that incorporating DNAm biomarkers from the original tissue can sometimes introduce noise to its prediction, with pure methylation loci offering clearer signals for certain biomarkers. DNAmCystatinC was one biomarker where the C method out performed the C+S method with a 0.259 improvement in correlation and 9 figure improvement in MSE compared to C+S method and saliva surrogate.

While not all TL algorithms outshined their corresponding saliva surrogates, they did exhibit substantial predictive power for 23 out of the 26 biomarkers. For instances where saliva DNAm biomarkers cannot be calculated without extensive imputation—as with the 40K array— our C method algorithms provide a valuable alternative. This resulted in 10 biomarkers having both a C+S algorithm and C algorithm. Three biomarkers have a C algorithm excluded due to inadequate predictive power: CD8pCD28nCD45RAn, Gran, and PlasmaBlast. In addition, we want to iterate that for CD4.naive, CD8.naive, DNAmB2M, and DNAmPAI1 biomarkers, the C algorithms are only recommended when saliva DNAm biomarkers are unavailable. A summary table detailing the available algorithms by biomarker can be found in Table 3.

**Table 3.** Summary of Cross Tissue DNAm Biomarker Algorithms Provided.

| Biomarker | Best Model | C Algorithm | Total Algorithms available |
| --- | --- | --- | --- |
| CD4.naive | Saliva | Yes | 1 |
| CD8.naive | Saliva | Yes | 1 |
| CD8pCD28nCD45RA<sub>n</sub> | Saliva | No | 0 |
| DNAmADM | C+S | Yes | 2 |
| DNAmAge | C | Yes | 1 |
| DNAmAgeHannum | C+S | Yes | 2 |
| DNAmB2M | Saliva | Yes | 1 |
| DNAmCystatinC | C | Yes | 1 |
| DNAmFEV1_noAge | C+S | Yes | 2 |
| DNAmFitAge | C+S | Yes | 2 |
| DNAmGait_noAge | C | Yes | 1 |
| DNAmGDF15 | C+S | Yes | 2 |
| DNAmGrimAge2BasedOnPredictedAge | C+S | Yes | 2 |
| DNAmGrimAge2BasedOnRealAge | C+S | Yes | 2 |
| DNAmGrimAgeBasedOnPredictedAge | C | Yes | 1 |
| DNAmGrimAgeBasedOnRealAge | C | Yes | 1 |
| DNAmGrip_noAge | C | Yes | 1 |
| DNAmLeptin | C+S | Yes | 2 |
| DNAmPACKYRS | C+S | Yes | 2 |
| DNAmPAI1 | Saliva | Yes | 1 |
| DNAmPhenoAge | C+S | Yes | 2 |
| DNAmTIMP1 | C | Yes | 1 |
| DNAmTL | C | Yes | 1 |
| DNAmVO2max | C | Yes | 1 |
| Gran | None | No | 0 |
| PlasmaBlast | C+S | No | 1 |

In summary, 11 biomarkers were best predicted using the C+S method, 9 with the C method, 5 using saliva DNAm biomarkers directly, and 1 biomarker was not well estimated by any method. We provide algorithms for predicting 23 blood DNAm biomarkers from saliva DNAm along with guidelines for their use. For researchers in the field, this study offers a comprehensive suite of algorithms tailored to a variety of biomarkers, enhancing the predictability and applicability of DNAm studies. Our research underscores the efficacy of TL in biomarker prediction and development and encourages its adoption over Lasso regression when applicable, particularly when multi-tissue DNAm data are available.

### 3.4 Application to Validation Datasets

#### 3.4.1 DNAmTL to Sex Relationship

We assessed the association between predicted blood DNA methylation Telomere Length (DNAmTL) biomarkers and sex, noting that females generally exhibit longer telomeres compared to males. Utilizing saliva DNA methylation (DNAm) as input, both C+S and C method blood DNAmTL predictions accurately reflected this trend. Specifically, females exhibited an average increase in telomere length of 0.29 and 0.3 for C+S and C methods, respectively, aligning closely with the observed 0.33 mean difference in the saliva surrogate. These disparities were statistically significant, as indicated by t-test and Kruskal Wallis test, with p-values ranging from 8.3E-6 to 3.5E-4 (Supplemental Table 2).

#### 3.4.2 COPD, Asthma, and Healthy Controls

Subsequently, we compared predicted blood DNAm biomarkers among individuals with Chronic Obstructive Pulmonary Disease (COPD), asthma, and healthy controls, adjusting for age using sputum DNAm as input. Consistent with previous studies showing advanced DNAm age in COPD and asthma patients, our analysis identified elevated sputum DNAm biomarkers in COPD individuals. Notably, a previously unreported association was discovered with DNAmFitAge, indicating that COPD patients have a 5.38 older sputum DNAmFitAge relative to healthy controls after age adjustment (p=0.036). For asthma, two C+S predicted biomarkers (DNAmPhenoAge and DNAmGDF15) aligned with expectations, whereas DNAmVO2max demonstrated an inverse relationship potentially influenced by inhaler use. For COPD patients, three predicted biomarkers corresponded with expected outcomes, revealing higher mean Smoking Pack Years (14.3, p=0.031) and an average 4.24 older DNAmAge (p=0.040) (Table 4A).

**Table 4.**
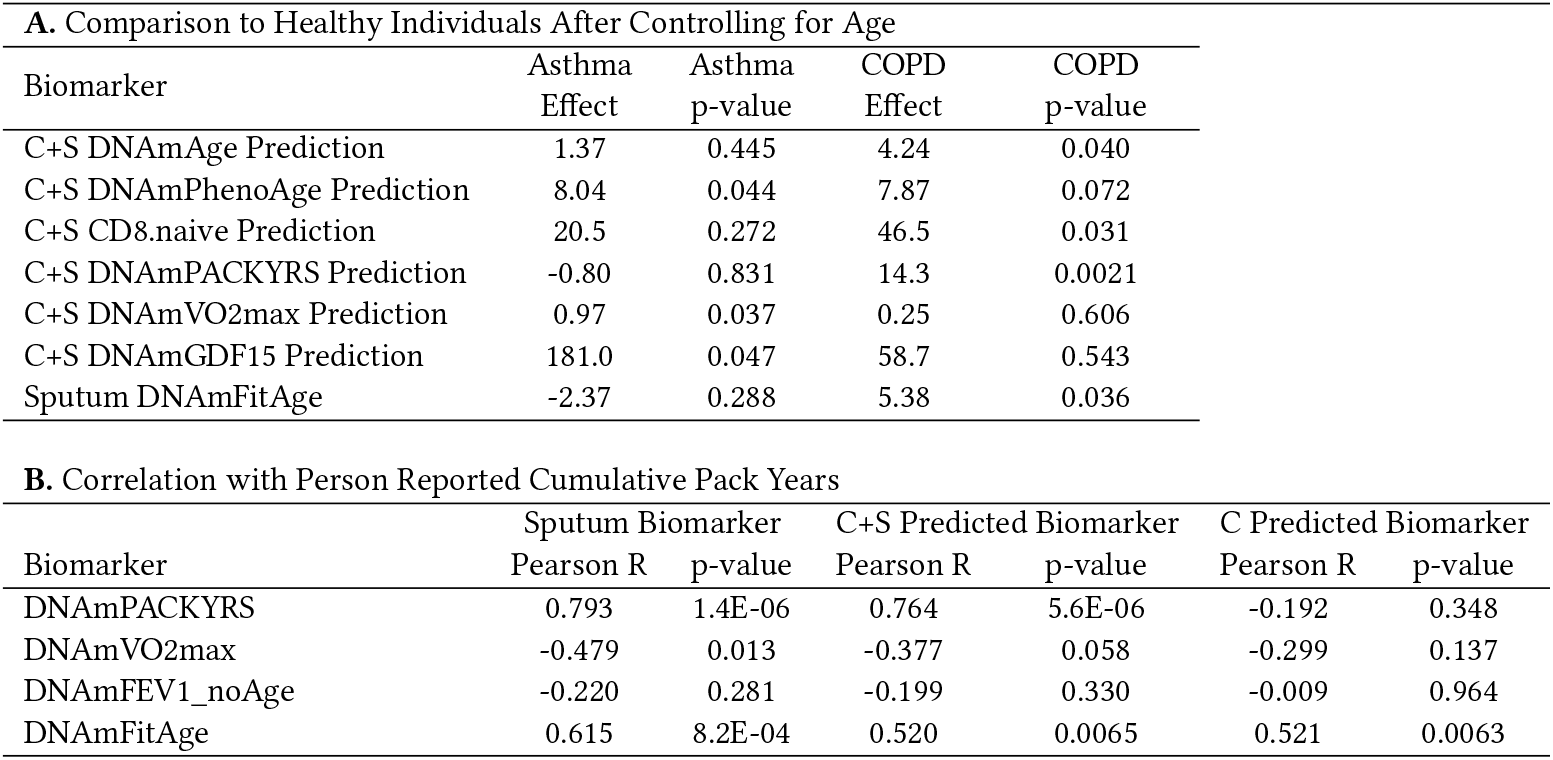
COPD, Asthma, and Healthy Control Analysis.

Our study further investigated the relationship between cumulative pack years and predicted DNAm biomarkers pertinent to lung health and fitness: DNAmPackYears, DNAmFEV1, and DNAmVO2max. Strong correlations were observed between cumulative pack years with sputum and blood predicted C+S DNAmPackYears: 0.793 (p=1.4E-6) and 0.764 (p=5.6E-6), respectively. However, the C DNAmPackYears did not significantly correlate with cumulative pack years (p=0.348), likely due to it being an inferior algorithm as demonstrated in the Leave-One-Out-Dataset (LODO) samples. Both sputum and blood-predicted C+S DNAmPackYears and DNAmVO2max exhibited the anticipated directional correlation with cumulative pack years, with DNAmVO2max inversely associated, suggesting diminished DNAmVO2max in longer-term smokers. Furthermore, both sputum and blood-predicted C+S and C DNAmFitAge displayed significant correlations with self-reported cumulative pack years, moving in the expected direction, with both C+S and C methods exhibiting comparable correlations of 0.52. These findings on DNAmVO2max and DNAmFitAge are novel, illustrating that health phenotypes associated with physical fitness and lifestyle factors like smoking can be detected using predicted blood DNAm biomarkers derived from methylation profiles of alternative tissues. This insight underscores the broader systemic implications of fitness and health habits, which manifest beyond the typically studied tissues, highlighting the potential for a more holistic understanding of health and disease (Table 4B).

#### 3.4.3 Exercise, Diet, and Sleep Intervention

Lastly, we evaluated the responsiveness of predicted blood DNAm biomarkers to a longitudinal 8-week intervention study focusing on an exercise, diet, and sleep. Employing a main effects mixed model, we controlled for age and study duration to ascertain if fitness-related blood biomarkers demonstrated anticipated improvements in the intervention group. This analysis provided a novel opportunity to assess the effectiveness of newly developed blood DNAm fitness biomarkers. We observed a significant increase in saliva DNAmVO2max by 0.781 in the intervention group (p=0.026), marking an unprecedented finding in exercise-induced DNAm fitness biomarker improvement in just 2 months time. Additionally, blood-predicted C DNAmCystatinC (a marker of kidney function), DNAmTL, DNAmVO2max, and DNAmFEV1_noAge exhibited significant enhancements in the intervention cohort. Notably, CystatinC decreased, while telomere length, VO2max, and FEV1 increased, with the latter two showing average improvements of 0.101 liters and 0.699 mL/kg/min, respectively. These changes paralleled the saliva DNAmVO2max improvement (0.781, p=0.026), reinforcing the credibility of our C algorithm for predicting DNAmVO2max (Table 5).

**Table 5.** Exercise, Diet, and Sleep Intervention Effects Controlling for Age, Study Time, and Person-Specific Variation.

| Outcome | Treatment Effect | p-value |
| --- | --- | --- |
| C DNAmCystatinC Prediction | -16580 | 0.030 |
| C DNAmTL Prediction | 0.112 | 0.031 |
| C DNAmFEV1_noAge Prediction | 0.101 | 0.033 |
| C DNAmVO2max Prediction | 0.699 | 0.033 |
| Saliva DNAmVO2max | 0.781 | 0.026 |

#### 3.4.4 Alternative Tissues

Because our TL approach incorporates information from multiple human tissues, we were interested in understanding if the algorithm is robust to tissue type. We applied the C algorithms to three accessible tissues (lymph nodes, adipose, and muscle DNAm) in HIV-negative and HIV-positive individuals and compared the predicted DNAm biomarkers. In lymph node DNAm samples, four predicted DNAm biomarkers demonstrated significant associations in the anticipated direction with HIV status after adjusting for age. Notably, DNAmAge and DNAmPhenoAge were, on average, 6.1 and 9.4 years older, respectively, in HIV-positive individuals when accounting for chronological age (p=0.043 and p=0.012), as shown in Table 6. These findings suggest that lymph nodes might serve as viable alternative tissue for our C algorithms as they can capture some biologically known differences. Nevertheless, the limited sample size (n=28) necessitates cautious interpretation, particularly concerning the performance of the remaining 19 predicted DNAm biomarkers using lymphatic tissue.

Conversely, adipose and muscle tissues exhibited markedly poor performance with these algorithms. Specifically, nine predicted biomarkers in adipose tissue and four in muscle tissue were significantly associated with HIV status after controlling for age. However, all 13 of these biomarkers displayed effects opposing the expected direction. Consequently, these outcomes imply that adipose and muscle tissues are unsuitable for application with our current C algorithms. The distinct responses across tissues underscore the necessity for a nuanced approach to selecting appropriate tissues for DNAm biomarker analysis in the context of HIV status and potentially other conditions (Table 6).

### 3.5 EpigenTL Software

Our methods and cross tissue prediction algorithms are freely available for researchers to use on Github at github.com/kristenmcgreevy/EpigenTL. Researchers interested in using transfer learning for their data can use function `Epigen.TL.Lasso`. To use this function, users will need to provide the matrix of covariates and outcome vector for the target and auxiliary datasets and specify the TL method to use. The methods are either of the three methods used to combine auxiliary datasets with the target data explored in this paper: Oracle A0, Oracle 1df, or Estimate A0. Other inputs can be specified by the user, including the coefficient threshold, number of Rhats to use, and how lambda is calculated. The function will return the transfer learning coefficients. We hope this function will be used by many to incorporate data from multiple sources, like directly incorporating animal data to improve development of new biomarkers with limited data, like those involved with brain aging.

While our methods were developed with cross tissue prediction for DNA methylation in mind, this methodology is not limited to methylation, and can be applied to any continuous outcome. We hope these methods will be used in expanding areas to develop other biomarkers not including methylation-based surrogate biomarkers.

For researchers interested in acquiring blood DNAm aging biomarkers by measuring saliva methylation instead of blood, they can use the function `Saliva.2.Blood.DNAmBiomarkers`. Users supply the saliva methylation values and specify whether to calculate biomarkers using the C or C+S algorithms. Because DNAmGrimAge is proprietary, algorithms for cross tissue prediction of DNAmGrimAge are excluded from online sources, however, researchers at a university can contact authors (KMM) directly to acquire these algorithms.

## 4 DISCUSSION

In this study, we explored the utility of Transfer Learning (TL) methodologies for predicting blood DNA methylation (DNAm) biomarkers from saliva DNAm, a critical advancement in the non-invasive assessment of epigenetic biomarkers. Our comprehensive approach not only sheds light on optimal TL application strategies but also provides researchers with tools to estimate 24 blood DNAm biomarkers from saliva, as well as functions to incorporate TL into their work. This research underscores the potential of TL to enhance the prediction accuracy and development of DNAm biomarkers.

Through extensive experimentation with different parameter settings and auxiliary data configurations, we identified optimal strategies for applying transfer learning in our context. This includes determining the best set of auxiliary data, the informative auxiliary set, and tuning the combination of model coefficients for improved DNAm biomarker prediction. Notably, our models demonstrated superior blood DNAm biomarker prediction accuracy from saliva compared to saliva surrogates and Lasso models, highlighting the effectiveness of TL in integrating diverse datasets and the utility of incorporating auxiliary data in high-dimensional settings.

A critical aspect of TL is the integration of auxiliary datasets, and our exploration into oracle and estimation-based methods for determining dataset informativeness addresses a broader challenge in TL: the need to leverage additional data without compromising or misdirecting the predictive focus of the model. Our comprehensive analysis, coupled with provided functions and recommendations, empowers researchers to employ our methods and final algorithms for cross-tissue DNAm biomarker prediction effectively. The Oracle 1df method, in particular, demonstrated good performance across both C+S and C methods. The divergence in optimal coefficient thresholding between the C+S and C methods suggests that the inclusion of saliva DNAm biomarkers requires a different approach to noise control compared to using CpGs alone.

The ability of TL predictions to reflect known biological trends validates the biological relevance of our algorithms. We demonstrate they reflect the longer telomeres in females and successfully differentiate between COPD, asthma, and healthy controls, with certain predicted biomarkers aligning with known disease characteristics. We show that predicted DNAm fitness biomarkers are responsive to an intervention study, demonstrating the promise of our algorithms for monitoring lifestyle changes and health interventions. This not only validates our algorithms but also illustrates their potential for novel discoveries, as evidenced by the identification of elevated sputum DNAmFitAge in COPD patients and increased saliva DNAmVO2max following an 8 week exercise intervention. These findings demonstrate recently developed fitness DNAm biomarkers can be measured in saliva, expanding the horizons of non-invasive health monitoring.

We provide 34 algorithms to estimate 24 blood DNAm biomarkers from saliva DNAm. We provide a comprehensive outline for preferred algorithm usage in various settings, and include 10 additional C algorithms with good predictive power for cases where saliva DNAm biomarkers are unmeasurable - like due to array differences. By developing our algorithms on CpG loci measured across multiple arrays, we ensure broad compatibility and utility.

The strength of our study lies in its extensive evaluation of TL algorithms across various parameters, coupled with comparisons to conventional methods. This provides a robust framework for understanding the relative performance of different approaches and offers detailed guidelines for employing TL in practice. The versatility and broader applicability of our algorithms are demonstrated through their successful application to various validation datasets, including alternative tissues. The ability of the algorithms to perform well in other tissues, like lymph nodes, provides promise for their application and reliability in other tissues by capturing shared information across different DNA methylation profiles. However, the observed poor performance in adipose and muscle tissues highlights the necessity of verifying biomarker reliability when applied to non-saliva tissues, emphasizing the need for careful tissue selection in research and clinical applications.

Our approach, rooted in the familiar terrain of penalized linear regression, aims to offer epigenetic and aging researchers advancements without overwhelming complexity. While the simplicity of linear regression broadens accessibility and improves its potential methodological integration, it does have limitations. For example, linear regression assumes a linear relationship between the saliva CpGs and the DNAm biomarker. Other methods, like Random forest and boosted tree models, may be better able to incorporate non-linear relationships, such as the presence of SNPs into models. Future research could explore TL frameworks incorporating non-linear or advanced regression models, potentially capturing complex biological relationships. Furthermore, our methods only employed 1 method for estimating auxiliary data informativeness, and future research could explore other methods not based in marginal associations.

These insights open new avenues for non-invasive biomarker development and facilitate crosstissue studies. The methodologies we propose are particularly advantageous for creating biomarkers in tissues that are traditionally challenging to access or available only in limited quantities, such as the brain and other internal organs. Additionally, given expansive data derived from animal models, we envision our outlined TL methods as ways for future research to integrate animal data as auxiliary datasets. The capacity of our approaches to accurately estimate informative datasets is a crucial asset in this context. It acts as a safeguard, ensuring that non-informative or distantly related data does not compromise the model’s integrity. This approach has the potential to significantly accelerate biomarker development, particularly for human tissues that are typically elusive to study. We strongly encourage researchers to adopt these TL strategies, not only as a means to expand the utility of extensive animal data but also to transform it into actionable insights and human biomarkers.

The study’s results provide valuable insights into the optimal parameters and comparative performance of TL algorithms in predicting blood DNAm biomarkers from saliva DNAm. The findings demonstrate the potential of TL to enhance the prediction accuracy of DNAm biomarkers, offer guidelines for selecting the right TL approach, and showcase the algorithms’ applicability to various biological and clinical scenarios. These contributions significantly advance the field of epigenetic research and open new avenues for non-invasive biomarker development and cross-tissue studies.

## 5 ACKNOWLEDGMENTS

## Notes

### Competing Interest Statement

The authors have declared no competing interest.

