## Supplemental Files for "Cross Tissue DNAm Biomarker Prediction using Transfer Learning"

### 6 SUPPLEMENTARY MATERIAL

**Supplemental Table 1.** Complete Comparison Among Transfer Learning, Lasso, and Saliva Surrogate Methods to Estimate Blood DNAm Biomarkers in C+S Methods

| | Transfer Learning C+S Method | | Transfer Learning C Method | | Lasso with Saliva DNAm Biomarkers | | Saliva DNAm Biomarkers | | C+S to Saliva | | C+S to Lasso | | C to C+S | | C to Saliva | |
|---|---|---|---|---|---|---|---|---|---|---|---|---|---|---|---|---|
| Biomarker | MSE | Corr | MSE | Corr | MSE | Corr | MSE | Corr | Corr Diff | MSE Diff | Corr Diff | MSE Diff | Corr Diff | MSE Diff | Corr Diff | MSE Diff |
| CD4.naive | 5.4E+04 | 0.285 | 4.5E+04 | 0.54 | 1.1E+06 | 0.597 | 4.7E+04 | 0.590 | -0.305 | -7413.94 | -0.312 | 1.1E+06 | 0.255 | 9887.86 | -0.050 | 2473.92 |
| CD8.naive | 9.3E+03 | 0.104 | 1.0E+04 | 0.274 | 5.9E+05 | -0.091 | 8.3E+03 | 0.480 | -0.376 | -962.02 | 0.195 | 5.8E+05 | 0.170 | -807.92 | -0.206 | -1769.95 |
| CD8pCD28nCD45RAn | 51.23 | -0.165 | 55.9 | -0.0831 | 525.5 | 0.118 | 52.7 | 0.335 | -0.500 | 1.50 | -0.283 | 474.2 | 0.081 | -4.67 | -0.418 | -3.17 |
| DNAmADM | 1198.0 | 0.641 | 2380.0 | 0.665 | 28482.6 | -0.368 | 2.1E+03 | 0.525 | 0.116 | 909.65 | 1.009 | 27284.7 | 0.024 | -1182.04 | 0.140 | -272.39 |
| DNAmAge | 75.31 | 0.873 | 75.4 | 0.937 | 1265.2 | 0.476 | 164.9 | 0.713 | 0.160 | 89.64 | 0.398 | 1189.9 | 0.064 | -0.09 | 0.224 | 89.55 |
| DNAmAgeHannum | 107.5 | 0.845 | 112.0 | 0.832 | 6391.1 | -0.128 | 227.9 | 0.703 | 0.141 | 120.38 | 0.972 | 6283.5 | -0.013 | -4.46 | 0.129 | 115.92 |
| DNAmB2M | 1.0E+11 | 0.627 | 1.1E+11 | 0.781 | 2.3E+11 | 0.732 | 5.0E+10 | 0.650 | -0.024 | -5.2E+10 | -0.106 | 1.3E+11 | 0.154 | -1.19E+10 | 0.131 | -6.4E+10 |
| DNAmCystatinC | 9.3E+09 | 0.415 | 5.6E+09 | 0.674 | 4.1E+10 | 0.638 | 8.7E+09 | 0.513 | -0.099 | -6.3E+08 | -0.223 | 3.2E+10 | 0.259 | 3.74E+09 | 0.161 | 3.1E+09 |
| DNAmFEV1_noAge | 0.47 | 0.588 | 0.65 | 0.589 | 14.50 | -0.261 | 0.6 | 0.630 | -0.042 | 0.10 | 0.849 | 14.0 | 0.001 | -0.18 | -0.041 | -0.08 |
| DNAmFitAge | 131.3 | 0.875 | 134.0 | 0.896 | 1894.0 | 0.399 | 211.5 | 0.721 | 0.154 | 80.13 | 0.476 | 1762.7 | 0.021 | -2.68 | 0.175 | 77.45 |
| DNAmGait_noAge | 0.05 | 0.803 | 0.0485 | 0.83 | 4.98 | 0.258 | 0.1 | 0.535 | 0.268 | 0.03 | 0.545 | 4.9 | 0.027 | 0.00 | 0.295 | 0.03 |
| DNAmGDF15 | 6.2E+04 | 0.661 | 1.1E+05 | 0.437 | 1.0E+06 | 0.024 | 6.8E+04 | 0.694 | -0.033 | 6477.09 | 0.638 | 9.6E+05 | -0.224 | -4.75E+04 | -0.257 | -4.1E+04 |
| DNAmGrimAge2Based OnPredictedAge | 151.9 | 0.775 | 207 | 0.689 | 2175.0 | -0.302 | 206.1 | 0.754 | 0.021 | 54.22 | 1.077 | 2023.1 | -0.086 | -55.10 | -0.065 | -0.88 |
| DNAmGrimAge2Based OnRealAge | 124.6 | 0.741 | 150 | 0.792 | 1292.1 | 0.235 | 196.1 | 0.694 | 0.048 | 71.58 | 0.506 | 1167.5 | 0.051 | -25.43 | 0.098 | 46.15 |
| DNAmGrimAgeBased OnPredictedAge | 168.8 | 0.749 | 109 | 0.881 | 876.2 | 0.157 | 141.0 | 0.791 | -0.043 | -27.89 | 0.592 | 707.3 | 0.132 | 59.84 | 0.090 | 31.95 |
| DNAmGrimAgeBased OnRealAge | 151.3 | 0.701 | 94.4 | 0.872 | 684.2 | 0.671 | 126.9 | 0.741 | -0.039 | -24.41 | 0.030 | 532.9 | 0.171 | 56.90 | 0.131 | 32.49 |
| DNAmGrip_noAge | 27.93 | 0.885 | 21.6 | 0.897 | 800.3 | 0.302 | 92.4 | 0.521 | 0.364 | 64.44 | 0.583 | 772.3 | 0.012 | 6.33 | 0.376 | 70.77 |
| DNAmLeptin | 3.7E+07 | 0.660 | 9.0E+07 | 0.425 | 2.3E+09 | -0.221 | 1.2E+08 | 0.302 | 0.359 | 8.5E+07 | 0.881 | 2.2E+09 | -0.235 | -5.32E+07 | 0.123 | 3.2E+07 |
| DNAmPACKYRS | 136.9 | 0.835 | 183 | 0.517 | 1049.7 | 0.774 | 125.3 | 0.690 | 0.145 | -11.68 | 0.061 | 912.7 | -0.318 | -46.05 | -0.173 | -57.73 |
| DNAmPAI1 | 2.5E+07 | 0.351 | 2.7E+07 | 0.503 | 1.1E+08 | -0.014 | 1.1E+07 | 0.450 | -0.098 | -1.4E+07 | 0.365 | 8.6E+07 | 0.152 | -1.81E+06 | 0.053 | -1.6E+07 |
| DNAmPhenoAge | 188.5 | 0.768 | 228 | 0.758 | 5612.1 | -0.192 | 306.6 | 0.718 | 0.050 | 118.09 | 0.959 | 5423.7 | -0.010 | -39.52 | 0.040 | 78.57 |
| DNAmTIMP1 | 4.6E+06 | 0.731 | 3.9E+06 | 0.905 | 7.0E+07 | 0.804 | 5.0E+06 | 0.724 | 0.006 | 4.4E+05 | -0.073 | 6.5E+07 | 0.174 | 7.31E+05 | 0.181 | 1.2E+06 |
| DNAmTL | 0.30 | 0.515 | 0.257 | 0.622 | 57.57 | -0.494 | 0.5 | 0.541 | -0.025 | 0.21 | 1.009 | 57.3 | 0.107 | 0.04 | 0.081 | 0.25 |
| DNAmVO2max | 7.36 | 0.625 | 7.04 | 0.736 | 2337.0 | 0.451 | 8.1 | 0.618 | 0.007 | 0.72 | 0.174 | 2329.6 | 0.111 | 0.32 | 0.118 | 1.04 |
| Gran | 0.13 | -0.109 | 0.137 | -0.232 | 1.42 | -0.390 | 0.2 | -0.208 | 0.099 | 0.05 | 0.281 | 1.3 | -0.123 | -0.01 | -0.024 | 0.04 |
| PlasmaBlast | 0.20 | 0.246 | 0.236 | 0.0297 | 3.12 | 0.017 | 0.4 | 0.010 | 0.236 | 0.19 | 0.229 | 2.9 | -0.216 | -0.04 | 0.020 | 0.15 |

**Supplemental Table 2.** Relationship of DNAmTL to Sex in GSE119078

| Biomarker | Mean Males (n=25) | Mean Females (n=34) | Females - Males | t-test p-value | Kruskal Wallis p-value |
|---|---|---|---|---|---|
| Saliva DNAmTL | 6.54 | 6.88 | 0.33 | 1.4E-04 | 4.2E-04 |
| C+S DNAmTL Blood Prediction | 6.86 | 7.16 | 0.29 | 8.3E-06 | 2.1E-05 |
| C DNAmTL Blood Prediction | 7.20 | 7.50 | 0.30 | 5.3E-04 | 3.5E-04 |

**Table 6.** Evaluation of Lymph, Adipose, and Muscle Tissue as TL Algorithm Input After Controlling for Age in HorvathHIV dataset

| Biomarker | Lymph Node (n=28) | | Adipose (n=58) | | Muscle (n=57) | |
|---|---|---|---|---|---|---|
| | HIV + Effect | p-value | HIV + Effect | p-value | HIV + Effect | p-value |
| CD4.naive C Prediction | -39.6 | 0.113 | -21.5 | 0.213 | 2.06 | 0.801 |
| CD8.naive C Prediction | -23.8 | 0.177 | 0.39 | 0.957 | -9.26 | 0.164 |
| DNAmADM C Prediction | 10.01 | 0.095 | -7.36 | 0.095 | -10.6 | 0.014 |
| DNAmAge C Prediction | 6.11 | 0.043 | -2.72 | 0.091 | -3.13 | 0.050 |
| DNAmAgeHannum C Prediction | 5.59 | 0.080 | -1.64 | 0.285 | -1.69 | 0.201 |
| DNAmB2M C Prediction | 65721 | 0.064 | -32378 | 0.131 | -34684 | 0.114 |
| DNAmCystatinC C Prediction | 46021 | 0.018 | -21372 | 0.021 | -9953 | 0.141 |
| DNAmFEV1_noAge C Prediction | -0.24 | 0.168 | 0.36 | 0.009 | 0.26 | 0.038 |
| DNAmFitAge C Prediction | 2.45 | 0.295 | -4.66 | 0.014 | -1.69 | 0.278 |
| DNAmGait_noAge C Prediction | 0.04 | 0.243 | 0.04 | 0.107 | 0.02 | 0.409 |
| DNAmGDF15 C Prediction | 265 | 0.039 | -28.48 | 0.453 | -68.5 | 0.177 |
| DNAmGrimAge2BasedOnPredictedAge C Prediction | 3.52 | 0.127 | -3.00 | 0.047 | -0.79 | 0.543 |
| DNAmGrimAge2BasedOnRealAge C Prediction | 3.92 | 0.071 | -2.63 | 0.042 | -0.45 | 0.702 |
| DNAmGrimAgeBasedOnPredictedAge C Prediction | 4.24 | 0.067 | -3.90 | 0.030 | -1.10 | 0.440 |
| DNAmGrimAgeBasedOnRealAge C Prediction | 3.66 | 0.120 | -3.15 | 0.019 | -0.94 | 0.442 |
| DNAmGrip_noAge C Prediction | -3.63 | 0.197 | 4.22 | 0.045 | 4.03 | 0.064 |
| DNAmLeptin C Prediction | 894 | 0.696 | -1318 | 0.189 | 414.1 | 0.775 |
| DNAmPACKYRS C Prediction | 6.59 | 0.298 | -0.54 | 0.750 | 0.36 | 0.847 |
| DNAmPAI1 C Prediction | 519 | 0.182 | 5.07 | 0.983 | -51.4 | 0.822 |
| DNAmPhenoAge C Prediction | 9.45 | 0.012 | -2.25 | 0.199 | -1.56 | 0.355 |
| DNAmTIMP1 C Prediction | 931 | 0.060 | -561.5 | 0.080 | -298 | 0.191 |
| DNAmTL C Prediction | -0.12 | 0.458 | 0.01 | 0.870 | -0.02 | 0.633 |
| DNAmVO2max C Prediction | -0.94 | 0.098 | 0.97 | 0.009 | 0.65 | 0.049 |

### 7 EPIGENTL FUNCTIONS

All of these functions can be found on Github with examples and more details on their use and function calls: https://github.com/kristenmcgreevy/EpigenTL. These functions are supplements to Section 3.2.

#### 7.1 `Epigen.TL.Lasso` Function

`Epigen.TL.Lasso` performs Transfer Learning Lasso allowing either the Auxiliary Dataset Information to be known (Oracle or Oracle 1df) or estimated in the process (Estimate A0).

As necessary input, X_matrix is the matrix of covariates (nxp) concatenated between the Target and all Auxiliary Datasets, in that order. n should therefore be n_0 + n_1 + ... + n_k. X_matrix needs to be a matrix and not a dataframe to allow for matrix multiplication to be carried out in the function. Y_vector is the outcome vector to develop the TL Lasso model to estimate. It should align with your X_matrix with Target and Auxiliary outcomes concatenated and be a nx1 vector. N_vector is a vector of number of observations in the Target and each Auxiliary datasets in the order they are concatenated. AuxInformation is either "Estimate A0", "Oracle", or "Oracle 1df" to signify the informative auxiliary datasets need estimated (EstA0) or they should be treated as known. If known, they can either be treated as equally informative and combined into 1 dataset (Oracle), or they can be treated individually (Oracle 1df). RhatCount is either "n0/3" or a value between 1, ..., p to specify how many marginal correlations to consider when calculating information. This should be set to NULL if AuxInformation is "Oracle" or "Oracle 1df". LambdaType is either "Constant" or

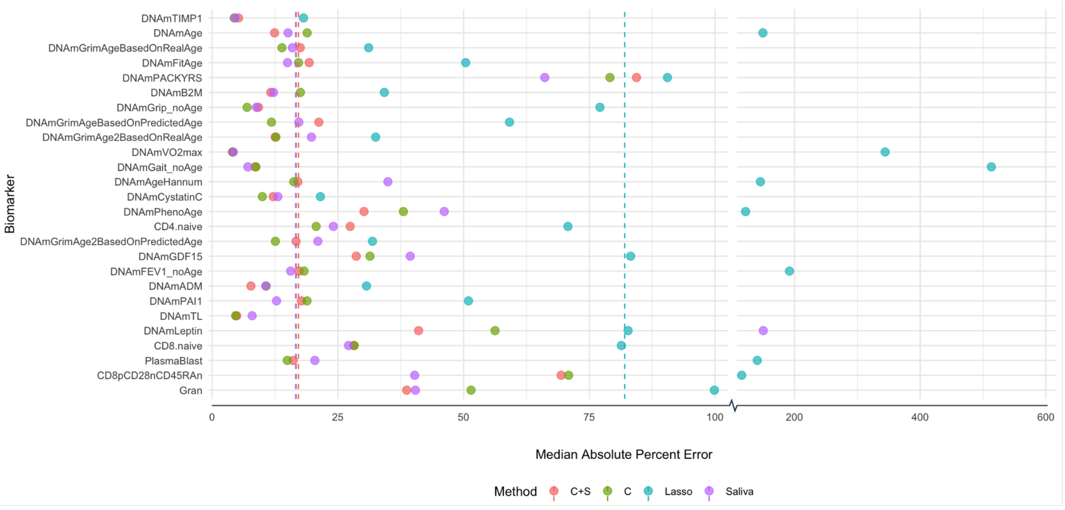


**Supplemental Figure 1.** Median Absolute Percent Error (MeAPE) between True and Estimated Blood DNAm Biomarkers by top Performing TL Methods, Lasso, and Saliva Surrogates. Median MeAPE presented as dotted line with a change in X axis scaling. LODO MeAPE presented for C+S, C, and Lasso methods.

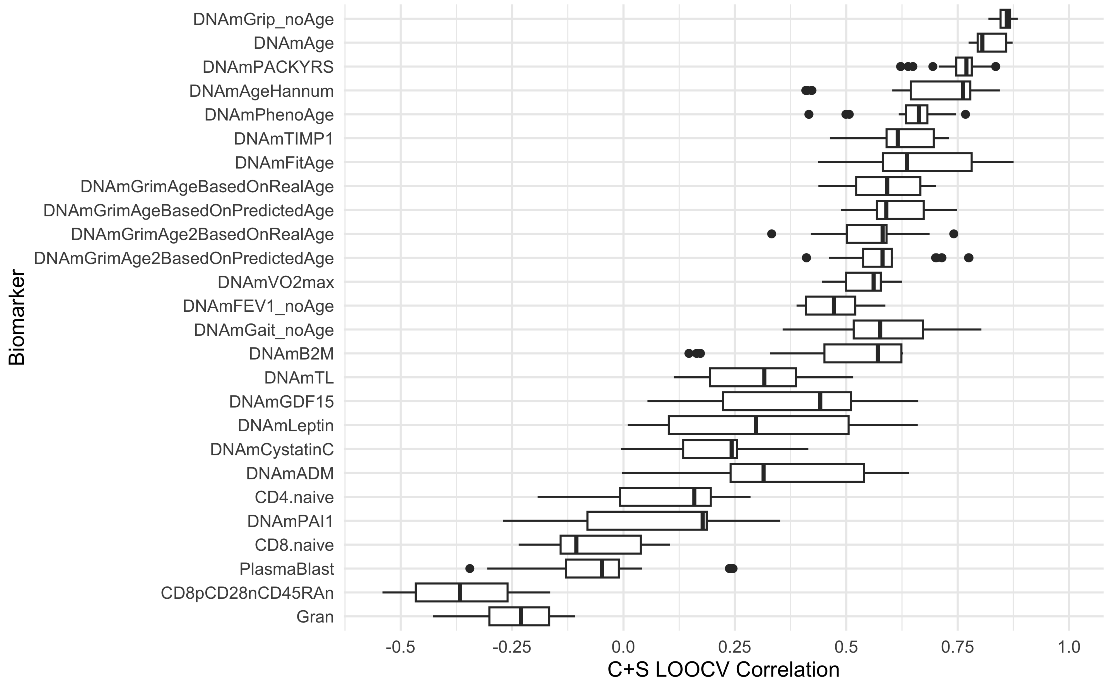


**Supplemental Figure 2.** Boxplots of C+S Method LODO Correlation by DNAm Biomarker to demonstrate variability in C+S method parameterizations.

"CV" to indicate if the lambda calculated from the target dataset should be used and adjusted based on the aux dataset size ("Constant") or whether the optimal lambda should be calculated via cross validation ("CV") for each set of auxiliary information. Default is "CV". seedstart is the seed to set in the calculation for reproducibility. If not specified, it is set to 123.

This function will return a data.frame with TL coefficients in the columns. The first column, "Variable" labels the intercept and columns from your X matrix that coefficient values correspond to. Columns 2:7 or 2:4 have the final coefficients with slight variations in their calculation. "min" and "1se" correspond to the lambda that either minimizes CV error or is at most 1se above it, respectively. "allcoef", "halfcoef", and "lambcoef" correspond to the coefficient thresholding used before combining coefficients across the auxiliary and target dataset. If "Constant" LambdaType was specified, only the minimum lambda from the target data is used and so "1se" coefficients are not presented.

```
Epigen.TL.Lasso <- function(X_matrix, Y_vector, N_vector,
                            AuxInformation, RhatCount = NULL, LambdaType = "CV",
                            seedstart = 123) {

  if(AuxInformation %!in% c("Estimate A0", "Oracle", "Oracle 1df")){
    stop("You must specify a method to capture auxiliary data informativeness.
         Options include 'Estimate A0', 'Oracle', or 'Oracle 1df'")
  }

  if(AuxInformation == "Estimate A0" & is.null(RhatCount)){
    stop("You must provide the number of Rhats to use when calculating
         auxiliary data informativeness.
        Options include any integer up to the column size of X_matrix OR 'n0/3'")
  }

  if(AuxInformation == "Estimate A0" & !is.null(RhatCount)){
    if(RhatCount > dim(X_matrix)[2]){
      stop("Rhat must be smaller than the number of columns in X.
        Please specify any integer up to the column size of X_matrix OR 'n0/3'")
    }

    if(RhatCount == 0){
     stop("Rhat must be a positive number from 1, ..., column size of X_matrix.")
    }
  }

  if(LambdaType %!in% c("CV", "Constant")){
    stop("LambdaType must be either 'CV' or 'Constant' to specify which
         lambda parameter is used for the auxiliary data.")
  }

  if(is.null(X_matrix)){
    stop("Please supply the X matrix")
  }
  if(is.null(Y_vector)){
```

```
    stop("Please supply the outcome")
  }
  if(is.null(N_vector)){
    stop("Please supply the N vector specifying the number of observations
         in target and auxiliary datasets")
  }

  if(sum(N_vector) != dim(X_matrix)[1]){
    stop("N_vector and X_matrix do not have the same number of observations")
  }

  if(sum(N_vector) != length(Y_vector)){
    stop("N_vector and Y do not have the same number of observations")
  }

  if(!is.null(RhatCount)){
    if(RhatCount == 'n0/3'){
        # set before calling the other functions.
        RhatCount <- round(N_vector[1] / 3)
    }
  }


    # TransLasso for Estimating informative set
    if(AuxInformation == "Estimate A0"){
      # for reproducibility
      set.seed(seedstart)

      Translasso_output <- TransLasso.EstA0(X = X_matrix, y = Y_vector,
                           n.vec = N_vector, AuxInformation = AuxInformation,
                           LambdaType = LambdaType, RhatCount = RhatCount)
    }
    # TransLasso for Oracle or Oracle 1df
    if(AuxInformation != "Estimate A0"){
      # for reproducibility
      set.seed(seedstart)

      Translasso_output <- TransLasso.Oracles(X = X_matrix, y = Y_vector,
                           n.vec = N_vector, AuxInformation = AuxInformation,
                           LambdaType = LambdaType)
    }

    # calculate intercepts
    intercept_beta_min <- mean(Y_vector - X_matrix %*%
                                 Translasso_output$beta.hat_min,
                               na.rm=TRUE)
    intercept_beta_1se <- mean(Y_vector - X_matrix %*%
                                 Translasso_output$beta.hat_1se,
```

```
                                    na.rm=TRUE)

    intercept_beta_min_lambda <- mean(Y_vector -
                        X_matrix %*% Translasso_output$beta.hat_min_lambda,
                                     na.rm=TRUE)
    intercept_beta_1se_lambda <- mean(Y_vector -
                        X_matrix %*% Translasso_output$beta.hat_1se_lambda,
                                     na.rm=TRUE)

    intercept_beta_min_halflambda <- mean(Y_vector -
                        X_matrix %*% Translasso_output$beta.hat_min_halflambda,
                                    na.rm=TRUE)
    intercept_beta_1se_halflambda <- mean(Y_vector -
                        X_matrix %*% Translasso_output$beta.hat_1se_halflambda,
                                    na.rm=TRUE)

    # keep all coefficients.
    TL_beta_coef <- data.frame(Variable = c("Intercept", colnames(X_matrix)),
                        beta_min_allcoef = c(intercept_beta_min,
                        Translasso_output$beta.hat_min),
                        beta_1se_allcoef = c(intercept_beta_1se,
                        Translasso_output$beta.hat_1se),
                        beta_min_halfcoef = c(intercept_beta_min_halflambda,
                        Translasso_output$beta.hat_min_halflambda),
                        beta_1se_halfcoef = c(intercept_beta_1se_halflambda,
                        Translasso_output$beta.hat_1se_halflambda),
                        beta_min_lambcoef = c(intercept_beta_min_lambda,
                        Translasso_output$beta.hat_min_lambda),
                        beta_1se_lambcoef = c(intercept_beta_1se_lambda,
                        Translasso_output$beta.hat_1se_lambda))

    # if Constant, only the min lambda is used.
    if(LambdaType == "Constant"){
        TL_beta_coef <- data.frame(Variable = c("Intercept", colnames(X_matrix)),
                             beta_min_allcoef = c(intercept_beta_min,
                                Translasso_output$beta.hat_min),
                            beta_min_halfcoef = c(intercept_beta_min_halflambda,
                                Translasso_output$beta.hat_min_halflambda),
                             beta_min_lambcoef = c(intercept_beta_min_lambda,
                                Translasso_output$beta.hat_min_lambda))
    }

  # return Final TL coefficients as a dataframe
  return(TL_beta_coef)

}
```

### 7.2 Saliva.2.Blood.DNAmBiomarkers Function

Saliva.2.Blood.DNAmBiomarkers functions calculates the blood DNAm Biomarkers from methylation values using either the C or C+S method algorithms.

As input, X is the matrix or dataframe of saliva DNA methylation beta values with samples in rows and methylation sites in columns. To see the list of CpG loci needed for computation, please call colnames to CS_Algorithms_GitHub or C_Algorithms_GitHub (which are loaded in the global environment). method should be either "C+S" or "C" to indicate which set of algorithms you are interested in calculating. If "C+S", you must also provide SalivaDNAmBiom. SalivaDNAmBiom should be the matrix or dataframe of Saliva DNAm biomarkers (ie DNAm biomarkers directly calculated with saliva methylation values) if using the C+S algorithms. Otherwise, this should be NULL. Default is NULL.

This function outputs a dataframe with predicted DNAm Biomarkers in each column with rows in the same order as the originally supplied X.

```
Saliva.2.Blood.DNAmBiomarkers <- function(X, method, SalivaDNAmBiom = NULL) {

  # make sure ppl provide the saliva DNAm biomarkers if its C+S
  if (method == "C+S" && is.null(SalivaDNAmBiom)) {
    stop("For method C+S, saliva DNAm Biomarker matrix must be provided.")
  }


  if (method == "C") {
    TLcoeffMatrix <- C_Algorithms_GitHub
    Varstart <- 2
  } else if (method == "C+S") {
    TLcoeffMatrix <- CS_Algorithms_GitHub
    Varstart <- 3
  } else {
    stop("Invalid method specified. Please specify either 'C' or 'C+S'")
  }


 cpgs_needed <- TLcoeffMatrix$Variable[Varstart:length(TLcoeffMatrix$Variable)]
  if(sum(cpgs_needed %!in% colnames(X)) > 0){
    cpgs_missing <<- cpgs_needed[cpgs_needed %!in% colnames(X)]
    stop("Not all CpG columns are present in X for this prediction.
         Please see cpgs_missing for a list of missing but necessary CpGs.")
  }


  X_keep <- X[, cpgs_needed]
  if (any(is.na(X_keep))) {
    stop("Missing values are not allowed.
            Please impute missing values in the X matrix.")
  }

  ### Now can start the computations ###
```

```
new_biomarker_list <- colnames(TLcoeffMatrix)[-1]

n_biom <- dim(TLcoeffMatrix)[2]
n_var <- dim(TLcoeffMatrix)[1]

# make temp dataset with intercept, saliva, and columns in correct order
if (method == "C"){
  newdata_temp <- data.frame(intercept = 1,
                             X[, TLcoeffMatrix$Variable[Varstart:n_var]])
}else{
  newdata_temp <- data.frame(intercept = 1, SalivaDNAmBiom = NA,
                             X[, TLcoeffMatrix$Variable[Varstart:n_var]])
}

# Turn Coefficient matrix into a real matrix for multiplying
TLcoeffMatrix2 <- matrix(unlist(TLcoeffMatrix[, 2:n_biom]),
                         ncol = (n_biom-1), nrow = n_var)

# initialize predictions dataframe
pred_df <- data.frame(rep(NA, dim(X)[1]))
k <- 1

for (i in new_biomarker_list) {

  if(method == "C+S"){
    # set Saliva value to the biomarker of interest
    newdata_temp$SalivaDNAmBiom <- SalivaDNAmBiom[, i]
  }

  # get the column of the TL coef matrix the biomarker is in
  TL_coef_Col <- (which(colnames(TLcoeffMatrix) == i) - 1)

  # turn into matrix for multiplication
  newdata_temp2 <- matrix(unlist(newdata_temp), ncol = dim(newdata_temp)[2],
                          nrow = dim(newdata_temp)[1])

  # data has ppl in rows, variables in columns,
  # TLcoef matrix has coefficients for each biomarker in a column
  cur_pred <- newdata_temp2 %*% TLcoeffMatrix2[, TL_coef_Col]

  # set prediction to the dataframe
  pred_df[, k] <- c(cur_pred)
  colnames(pred_df)[k] <- i

  # make k go up
  k <- k + 1
```

```
    # remove values so we don't repeat by accident
    rm(cur_pred); rm(newdata_temp2)

    if(method == "C+S"){
      # set Saliva value to NA so we don't repeat by accident
      newdata_temp$SalivaDNAmBiom <- NA
    }

  } # end of for loop in biomarker list

  # return the predicted DNAm values.
  if(method == "C+S"){
    colnames(pred_df) <- paste0(colnames(pred_df), "_CS_Pred")
  } else{
    colnames(pred_df) <- paste0(colnames(pred_df), "_C_Pred")
  }

  return(pred_df)
}
```

#### 7.3 TL_Lasso Function

`TL_Lasso` is the actual Transfer Learning Lasso loop process called inside both `TL.Lasso.EstA0` and `TL.Lasso.Oracles`. As input, X is the matrix of covariates, y is the outcome vector, A0 is the informative Aux Set. It is either NULL or estimated. n.vec is a vector of number of observations in the target and aux datasets in that order, lam.const is whether we are calculating the optimal lambda via CV in each informative aux set or the constant value if using the methods outlined in TransLasso paper.

It outputs the estimated beta coefficients with and without thresholding (when performed in auxiliary data) and the estimated lambda values.

```
TL_Lasso <- function(X, y, A0, n.vec, lam.const=NULL, lam.const_1se = NULL, ...){

  p <- ncol(X)
  size.A0 <- length(A0) # set to NULL so its 0

  if(size.A0 > 0){ # only for Aux data, otherwise SKIP to below

    ind.kA <- ind.set(n.vec, c(1, A0+1))
    ind.1 <- 1:n.vec[1] # vector of all values to build initial model.

    y.A <- y[ind.kA]

    # if null, CV done for each Informative Set and both min and 1se lambda kept.
    if(is.null(lam.const)){
    # this gets run on first run because we have it set to NULL
      # does its own grid search for lambda
      cv.init<-cv.glmnet(X[ind.kA,], y.A, nfolds=8)
    # now, it will just take whatever the best value was and calculate the constant.
```

```
    lam.const <- cv.init$lambda.min/sqrt(2*log(p)/length(ind.kA))
    lam.const_1se <- cv.init$lambda.1se/sqrt(2*log(p)/length(ind.kA))
  }
  if(!is.null(lam.const) & is.null(lam.const_1se)){
    lam.const_1se <- lam.const
  }

  # w.kA = coefficients from Xk predicts Yk
  w.kA_min <- as.numeric(glmnet(X[ind.kA,], y.A,
                  lambda=lam.const*sqrt(2*log(p)/length(ind.kA)))$beta)
  w.kA_1se <- as.numeric(glmnet(X[ind.kA,], y.A,
                  lambda=lam.const_1se*sqrt(2*log(p)/length(ind.kA)))$beta)

  # w.k coefficient thresholding
  w.kA_min_halflambda <- w.kA_min*(abs(w.kA_min) >=
                          0.5*lam.const*sqrt(2*log(p)/length(ind.kA)))
  w.kA_min_lambda <- w.kA_min*(abs(w.kA_min) >=
                           lam.const*sqrt(2*log(p)/length(ind.kA)))

  w.kA_1se_halflambda <- w.kA_1se*(abs(w.kA_1se) >=
                          0.5*lam.const_1se*sqrt(2*log(p)/length(ind.kA)))
  w.kA_1se_lambda <- w.kA_1se*(abs(w.kA_1se) >=
                          lam.const_1se*sqrt(2*log(p)/length(ind.kA)))

  # build model in target where outcome is what is left after
  # taking the yhat from kth model coefficients.
  # delta.kA = coefficients from X0 predicts (Y0 - Y0hat from w.kA)
 delta.kA_min <- as.numeric(glmnet(x=X[ind.1,],y=y[ind.1]-X[ind.1,]%*%w.kA_min,
                        lambda=lam.const*sqrt(2*log(p)/length(ind.1)))$beta)
 delta.kA_1se <- as.numeric(glmnet(x=X[ind.1,],y=y[ind.1]-X[ind.1,]%*%w.kA_1se,
                     lambda=lam.const_1se*sqrt(2*log(p)/length(ind.1)))$beta)

  # delta.k coefficient thresholding
  delta.kA_min_halflambda <- delta.kA_min*(abs(delta.kA_min) >=
                              0.5*lam.const*sqrt(2*log(p)/length(ind.1)))
  delta.kA_min_lambda <- delta.kA_min*(abs(delta.kA_min) >=
                              lam.const*sqrt(2*log(p)/length(ind.1)))

  delta.kA_1se_halflambda <- delta.kA_1se*(abs(delta.kA_1se) >=
                              0.5*lam.const_1se*sqrt(2*log(p)/length(ind.1)))
  delta.kA_1se_lambda <- delta.kA_1se*(abs(delta.kA_1se) >=
                              lam.const_1se*sqrt(2*log(p)/length(ind.1)))

  # final beta coefficients (the kth lasso coefficients are the weights
  # from kth lasso model + kth lasso on target)

  # no thresholding
  beta.kA_min <- w.kA_min + delta.kA_min
```

```
    beta.kA_1se <- w.kA_1se + delta.kA_1se

    # half lambda thresholding
    beta.kA_min_halflambda <- w.kA_min_halflambda + delta.kA_min_halflambda
    beta.kA_min_lambda <- w.kA_min_lambda + delta.kA_min_lambda

    # lambda thresholding
    beta.kA_1se_halflambda <- w.kA_1se_halflambda + delta.kA_1se_halflambda
    beta.kA_1se_lambda <- w.kA_1se_lambda + delta.kA_1se_lambda

   lam.const=NULL # reset lambda because we don't want it to loop through as !null
    # first auxillary dataset because we supply the lambda constant.

    # output all coefficients
    # recall if constant lambda was specified, min and 1se will be identical.
    list(beta.kA_min = as.numeric(beta.kA_min), w.kA_min=w.kA_min,
       beta.kA_1se = as.numeric(beta.kA_1se), w.kA_1se=w.kA_1se,
       beta.kA_min_halflambda = as.numeric(beta.kA_min_halflambda),
            w.kA_min_halflambda=w.kA_min_halflambda,
       beta.kA_1se_halflambda = as.numeric(beta.kA_1se_halflambda),
            w.kA_1se_halflambda=w.kA_1se_halflambda,
       beta.kA_min_lambda = as.numeric(beta.kA_min_lambda),
            w.kA_min_lambda=w.kA_min_lambda,
       beta.kA_1se_lambda = as.numeric(beta.kA_1se_lambda),
            w.kA_1se_lambda=w.kA_1se_lambda,
       lam.const=lam.const, lam.const_1se=lam.const_1se)
  }else{ # end of if(size.A0 > 0)

    # BEGINNING CODE FOR INITIAL / TARGET DATA
    cv.init <- cv.glmnet(X[1:n.vec[1],], y[1:n.vec[1]], nfolds=8)

    # When constant lambda selected, min lambda is used.
    lam.const <- cv.init$lambda.min/sqrt(2*log(p)/n.vec[1])
    lam.const_1se <- cv.init$lambda.1se/sqrt(2*log(p)/n.vec[1])

    # extract coefficients (excluding the intercept)
    beta.kA_min <- predict(cv.init, s='lambda.min', type='coefficients')[-1]
    beta.kA_1se <- predict(cv.init, s='lambda.1se', type='coefficients')[-1]
    w.kA_min <- w.kA_1se <- NA

    list(beta.kA_min = as.numeric(beta.kA_min), w.kA_min=w.kA_min,
       beta.kA_1se = as.numeric(beta.kA_1se), w.kA_1se=w.kA_1se,
       lam.const=lam.const, lam.const_1se = lam.const_1se)
  }

}
```

#### 7.4 TransLasso.Oracles Function

TransLasso.Oracles performs Oracle Transfer Learning Lasso meaning the set of auxiliary datasets is specified to either run all at once (Oracle) or 1 dataset at a time (Oracle 1df). As input, X is the matrix of covariates, y is the outcome vector, n.vec is a vector of number of observations in the target and aux datasets in that order, AuxInformation is either "Oracle" or "Oracle 1df", LambdaType is either "Constant" or "CV" to indicate if the lambda calculated from the target dataset should be used and adjusted based on the aux dataset size ("Constant") or whether the optimal lambda should be calculated via cross validation ("CV") for each set of auxiliary information.

It outputs the aggregated coefficients at various parameterizations and the weights used when aggregating.

```
TransLasso.Oracles <- function(X, y, n.vec, AuxInformation = "Oracle 1df",
                               LambdaType = "CV", ...) {

  M = length(n.vec)-1
  p <- ncol(X)

  # row indices of where these observations are for target
  ind.1 <- ind.set(n.vec, 1)

  Tset <- list()

  if(AuxInformation == "Oracle"){
      # make Tset actually just all the aux datasets (for oracle all at once)
      # aux datasets start at 2.
      Tset[[1]] <- c(1:M)
  }
  # take 1 aux dataset at a time, noting that we index by dataset before.
  if(AuxInformation == "Oracle 1df"){
    for(kk in 1:M){ #use Rhat as the selection rule
     Tset[[kk]] <- kk
    } # the sets of aux datasets to take for ranking of datasets.
  }

  k0 = length(Tset)
  Tset <- unique(Tset)

  beta.T_min <- beta.T_min_lambda <- beta.T_min_halflambda <- list()
  beta.T_1se <- beta.T_1se_lambda <- beta.T_1se_halflambda <- list()

  # Lasso on Target Data only
  init.re <- TL_Lasso(X=X, y=y, A0=NULL, n.vec=n.vec)

  beta.T_min[[1]] <- init.re$beta.kA_min
  beta.T_1se[[1]] <- init.re$beta.kA_1se

  # if constant lambda specified, it is here.
  c1_lambda_const <- init.re$lam.const
```

```
c1_lambda_const_1se <- init.re$lam.const_1se

og_lasso_coef_min <- init.re$beta.kA_min
og_lasso_coef_1se <- init.re$beta.kA_1se


beta.T_min_lambda <- beta.T_min_halflambda <- beta.T_min
beta.T_1se_lambda <- beta.T_1se_halflambda <- beta.T_1se

# go through TL Lasso for each informative set
for(kk in 1:length(Tset)){
  T.k <- Tset[[kk]]

  # changed function call to lam.const = NULL to do Aux CV
  if(LambdaType == "CV"){
    re.k <- TL_Lasso(X=X, y=y, A0=T.k, n.vec=n.vec, lam.const = NULL)
  }else{ # constant lambda specification
   re.k <- TL_Lasso(X=X, y=y, A0=T.k, n.vec=n.vec, lam.const = c1_lambda_const,
                    lam.const_1se = c1_lambda_const_1se)
  }

  # extract coefficients for each informative auxiliary set
  beta.T_min[[kk+1]] <- re.k$beta.kA_min
  beta.pool.T_min[[kk+1]] <- re.k$w.kA_min

  beta.T_1se[[kk+1]] <- re.k$beta.kA_1se
  beta.pool.T_1se[[kk+1]] <- re.k$w.kA_1se

  beta.T_min_halflambda[[kk+1]] <- re.k$beta.kA_min_halflambda
  beta.pool.T_min_halflambda[[kk+1]] <- re.k$w.kA_min_halflambda

  beta.T_1se_lambda[[kk+1]] <- re.k$beta.kA_1se_lambda
  beta.pool.T_1se_lambda[[kk+1]] <- re.k$w.kA_1se_lambda
}

beta.T_min <- beta.T_min[!duplicated((beta.T_min))]
beta.T_min <- as.matrix(as.data.frame(beta.T_min))

beta.T_1se <- beta.T_1se[!duplicated((beta.T_1se))]
beta.T_1se <- as.matrix(as.data.frame(beta.T_1se))

beta.T_min_halflambda <- beta.T_min_halflambda[
                          !duplicated((beta.T_min_halflambda))]
beta.T_min_halflambda <- as.matrix(as.data.frame(beta.T_min_halflambda))
beta.T_min_lambda <- beta.T_min_lambda[!duplicated((beta.T_min_lambda))]
beta.T_min_lambda <- as.matrix(as.data.frame(beta.T_min_lambda))

beta.T_1se_halflambda <- beta.T_1se_halflambda[
```

```
                              !duplicated((beta.T_1se_halflambda))]
  beta.T_1se_halflambda <- as.matrix(as.data.frame(beta.T_1se_halflambda))
  beta.T_1se_lambda <- beta.T_1se_lambda[!duplicated((beta.T_1se_lambda))]
  beta.T_1se_lambda <- as.matrix(as.data.frame(beta.T_1se_lambda))

  ## aggregate coefficients using squared error.
  ## aggregate w.kA and delta.kA
  agg.re1_min <- coef.aggr(B= beta.T_min, X = X, y = y, N_vector = n.vec)
  agg.re1_1se <- coef.aggr(B= beta.T_1se, X = X, y = y, N_vector = n.vec)

  agg.re1_min_halflambda <- coef.aggr(B= beta.T_min_halflambda, X = X, y = y,
                                      N_vector = n.vec)
  agg.re1_1se_halflambda <- coef.aggr(B= beta.T_1se_halflambda, X = X, y = y,
                                      N_vector = n.vec)

  agg.re1_min_lambda <- coef.aggr(B= beta.T_min_lambda, X = X,
                                  y = y, N_vector = n.vec)
  agg.re1_1se_lambda <- coef.aggr(B= beta.T_1se_lambda, X = X,
                                  y = y, N_vector = n.vec)



  return(list(beta.hat_min = agg.re1_min$beta, theta.hat_min = agg.re1_min$theta,
              beta.hat_1se = agg.re1_1se$beta, theta.hat_1se = agg.re1_1se$theta,
               beta.hat_min_halflambda = agg.re1_min_halflambda$beta,
               theta.hat_min_halflambda = agg.re1_min_halflambda$theta,
               beta.hat_1se_halflambda = agg.re1_1se_halflambda$beta,
               theta.hat_1se_halflambda = agg.re1_1se_halflambda$theta,
               beta.hat_min_lambda = agg.re1_min_lambda$beta,
               theta.hat_min_lambda = agg.re1_min_lambda$theta,
               beta.hat_1se_lambda = agg.re1_1se_lambda$beta,
               theta.hat_1se_lambda = agg.re1_1se_lambda$theta,
               c1_lambda_const = c1_lambda_const))
}
```

#### 7.5 `TransLasso.EstA0` Function

`TransLasso.EstA0` performs Transfer Learning Lasso while estimating the Informative Auxiliary Data. X is the matrix of covariates (nxp), y is the outcome vector (nx1), n.vec is a vector of number of observations in the target and aux datasets in that order, RhatCount is either "n0/3" or a value between 1, ..., p to specify how many marginal correlations to consider when calculating information. LambdaType is either "Constant" or "CV" to indicate if the lambda calculated from the target dataset should be used and adjusted based on the aux dataset size ("Constant") or whether the optimal lambda should be calculated via cross validation ("CV") for each set of auxiliary information.

It outputs the aggregated coefficients at various parameterizations and the weights used when aggregating.

```
TransLasso.EstA0 <- function(X, y, n.vec, RhatCount, LambdaType, ...){
```

```
# count of aux datasets
M = length(n.vec)-1

Rhat <- rep(0, M+1)
p <- ncol(X)

# make row indices of where target observations are
ind.1 <- ind.set(n.vec, 1)

# calculate informativeness for each aux study
for(k in 2: (M+1)){
  ind.k <- ind.set(n.vec, k) # row indices for kth aux sample.

  # calculate difference in marginal correlations between k aux and target data.
 Xty.k <- t(X[ind.k, ])%*%y[ind.k] / n.vec[k] - t(X[ind.1,])%*%y[ind.1]/ n.vec[1]

  # Rhat adjusts how many marginal correlations are looked at
  # take the top largest correlation differences based on RhatCount
  margin.T <- sort(abs(Xty.k), decreasing=T)[1:RhatCount]

  # estimated sparse index for kth aux sample.
  Rhat[k] <- sum(margin.T^2)
}

Tset <- list()
k0 = 0
# get ordering of smallest to largest Rhat for aux samples.
kk.list <- unique(rank(Rhat[-1]))

for(kk in 1:length(kk.list)){#use Rhat as the selection rule
  Tset[[k0+kk]] <- which(rank(Rhat[-1]) <= kk.list[kk])
} # the sets of aux datasets to take for each ranking of datasets to include.

k0 = length(Tset)
Tset <- unique(Tset)

beta.T_min <- beta.T_min_lambda <- beta.T_min_halflambda <- list()
beta.T_1se <- beta.T_1se_lambda <- beta.T_1se_halflambda <- list()

# Lasso on Target Data only
init.re <- TL_Lasso(X=X, y=y, A0=NULL, n.vec=n.vec)

beta.T_min[[1]] <- init.re$beta.kA_min
beta.T_1se[[1]] <- init.re$beta.kA_1se

# if constant lambda specified, it is here.
c1_lambda_const <- init.re$lam.const
```

```
c1_lambda_const_1se <- init.re$lam.const_1se

og_lasso_coef_min <- init.re$beta.kA_min
og_lasso_coef_1se <- init.re$beta.kA_1se

beta.T_min_lambda <- beta.T_min_halflambda <- beta.T_min
beta.T_1se_lambda <- beta.T_1se_halflambda <- beta.T_1se

# go through TL Lasso for each informative set
for(kk in 1:length(Tset)){
  T.k <- Tset[[kk]]

  # which lambda type changes which section of function call it goes into
  if(LambdaType == "CV"){
    re.k <- TL_Lasso(X=X, y=y, A0=T.k, n.vec=n.vec, lam.const = NULL)
  }
  if(LambdaType == "Constant"){
    re.k <- TL_Lasso(X=X, y=y, A0=T.k, n.vec=n.vec,
                     lam.const = c1_lambda_const,
                     lam.const_1se = c1_lambda_const_1se)
  }

  # extract coefficients for each informative auxiliary set
  beta.T_min[[kk+1]] <- re.k$beta.kA_min
  beta.T_1se[[kk+1]] <- re.k$beta.kA_1se
  beta.T_min_halflambda[[kk+1]] <- re.k$beta.kA_min_halflambda
  beta.T_min_lambda[[kk+1]] <- re.k$beta.kA_min_lambda
  beta.T_1se_lambda[[kk+1]] <- re.k$beta.kA_1se_lambda
  beta.T_1se_halflambda[[kk+1]] <- re.k$beta.kA_1se_halflambda
}

beta.T_min <- beta.T_min[!duplicated((beta.T_min))]
beta.T_min <- as.matrix(as.data.frame(beta.T_min))

beta.T_1se <- beta.T_1se[!duplicated((beta.T_1se))]
beta.T_1se <- as.matrix(as.data.frame(beta.T_1se))

beta.T_min_halflambda <- beta.T_min_halflambda[
                         !duplicated((beta.T_min_halflambda))]
beta.T_min_halflambda <- as.matrix(as.data.frame(beta.T_min_halflambda))
beta.T_min_lambda <- beta.T_min_lambda[!duplicated((beta.T_min_lambda))]
beta.T_min_lambda <- as.matrix(as.data.frame(beta.T_min_lambda))

beta.T_1se_halflambda <- beta.T_1se_halflambda[
                      !duplicated((beta.T_1se_halflambda))]
beta.T_1se_halflambda <- as.matrix(as.data.frame(beta.T_1se_halflambda))
beta.T_1se_lambda <- beta.T_1se_lambda[!duplicated((beta.T_1se_lambda))]
beta.T_1se_lambda <- as.matrix(as.data.frame(beta.T_1se_lambda))
```

```
  ## aggregate coefficients using squared error.
  # No coef thresholding
  agg.re1_min <- coef.aggr(B= beta.T_min, X = X, y = y, N_vector = n.vec)
  agg.re1_1se <- coef.aggr(B= beta.T_1se, X = X, y = y, N_vector = n.vec)

  # half lambda coef thresholding
  agg.re1_min_halflambda <- coef.aggr(B= beta.T_min_halflambda, X = X, y = y,
                                    N_vector = n.vec)
  agg.re1_1se_halflambda <- coef.aggr(B= beta.T_1se_halflambda, X = X, y = y,
                                    N_vector = n.vec)

  # lambda coef thresholding
  agg.re1_min_lambda <- coef.aggr(B= beta.T_min_lambda, X = X,
                  y = y, N_vector = n.vec)
  agg.re1_1se_lambda <- coef.aggr(B= beta.T_1se_lambda, X = X,
                  y = y, N_vector = n.vec)


  # theta are the returned weights of each dataset.
  # betas are the final, aggregated coefficients for potential covariates
 return(list(beta.hat_min = agg.re1_min$beta, theta.hat_min = agg.re1_min$theta,
            beta.hat_1se = agg.re1_1se$beta, theta.hat_1se = agg.re1_1se$theta,
             beta.hat_min_halflambda = agg.re1_min_halflambda$beta,
             theta.hat_min_halflambda = agg.re1_min_halflambda$theta,
             beta.hat_1se_halflambda = agg.re1_1se_halflambda$beta,
             theta.hat_1se_halflambda = agg.re1_1se_halflambda$theta,
             beta.hat_min_lambda = agg.re1_min_lambda$beta,
             theta.hat_min_lambda = agg.re1_min_lambda$theta,
             beta.hat_1se_lambda = agg.re1_1se_lambda$beta,
             theta.hat_1se_lambda = agg.re1_1se_lambda$theta,
             c1_lambda_const = c1_lambda_const,
             og_lasso_coef_min = og_lasso_coef_min,
             og_lasso_coef_1se = og_lasso_coef_1se))
}
```

#### 7.6 Helper Functions

`coef.aggr` function aggregates the coefficients from target and auxiliary data using the error in the target data. This function is called in the background; it is not being called directly by users.

As input, this function requires B, the coefficient vector, X, the matrix of covariates, y, the outcome vector, and N_vector, a vector of number of observations in the target and aux datasets in that order. It outputs a two item list, with the first item, theta, being the weights used to aggregate the coefficients, and beta, the final, aggregated coefficients.

```
coef.aggr <- function(B, X, y, N_vector){

  # if all coefficients are zero, just return zero
  if(sum(B == 0) == ncol(B)*nrow(B)){
```

```
    return(rep(0,nrow(B)))
  }
  p <- nrow(B)
  K <- ncol(B)
  colnames(B) <- NULL

    # Take the difference in target y and predicted y
    # from each aux data coefficients
    y0hatk <- -log(colSums(y[1:N_vector[1]] - X[1:N_vector[1], ] %*% B)^2)
    theta.hat <- exp(y0hatk)
    theta.hat = theta.hat / sum(theta.hat)
    # weights by fraction of total squared error

    # multiply betak by weights for each kth aux
    beta <- as.numeric(B%*%theta.hat)

  list(theta = theta.hat, beta = beta)

}
```

`ind.set` tells the TL functions where the first and last observation is for each dataset (target, aux 1, ..., aux k) based on the sample sizes in n.vec. n.vec is the vector of number of observations in the target and aux datasets in that order. k is index of the dataset we are interested in extracting values from. It returns the indices in the X matrix and y vector to extract.

```
ind.set <- function(n.vec, k.vec){
  ind.re <- NULL
  for(k in k.vec){
    if(k==1){
      ind.re<-c(ind.re,1: n.vec[1])
    }else{
      ind.re<- c(ind.re, (sum(n.vec[1:(k-1)])+1): sum(n.vec[1:k]))
    }
  }
  ind.re
}
```

Received February 2024
